# STRESS HISTORY ESTABLISHES A TRANSIENT TOLERANT STATE THAT SHAPES ANTIBIOTIC SURVIVAL UPON RESUSCITATION

**DOI:** 10.64898/2026.03.01.708845

**Authors:** Kieran Abbott, Georgeos Hardo, Ruizhe Li, Jack Bradley, Ashraf Zarkan, Somenath Bakshi

**Affiliations:** Department of Engineering, University of Cambridge; Department of Genetics, University of Cambridge; Department of Biology, United Arab Emirates University

## Abstract

Antibiotic treatment failure, often driven by non-genetic mechanisms such as tolerance and persistence, remains a major global health challenge. *β*-lactams, the most widely prescribed antibiotic class, are particularly compromised by tolerance in dormant, non-growing cells; yet, how these drugs act on cells resuscitating from dormancy remains poorly understood. Here, we investigate the resuscitation phase at an unprecedented scale using Hi-DFA (High-throughput Dynamic Fate Analyser), a single-cell microfluidic platform integrating time-lapse imaging with machine-learning-based image analysis for dynamic cell-fate tracking. We identify a distinct survival strategy: a significant fraction of resuscitating cells transiently slow their growth, facilitating survival upon *β*-lactam exposure. This ‘transiently tolerant’ phenotype is considerably less frequent in unstressed, exponentially growing cells, indicating that prior starvation history predisposes cells to this state. Using simulated *in vitro* pharmacokinetic treatment profiles, we show that suboptimal dosing selectively enriches for this transient tolerance state. A population dynamics model built from this single-cell antibiotic-response data suggests that these transient-tolerant cells, not typical starvation-triggered persisters, may be the primary drivers of rapid population regrowth post-treatment under clinically relevant conditions. Together, our findings define a distinct class of antibiotic survival shaped by stress history and treatment profile, offering a quantitative framework for optimising antibiotic dosing strategy.

## 1 Introduction

Antimicrobial resistance (AMR) remains one of the greatest threats to global health, responsible for over one million deaths each year[1]. Yet resistant infections account for less than 15% of the roughly nine million annual deaths attributed to bacterial disease[2, 3], underscoring that treatment failure often stems from factors beyond genetic resistance. While limited antibiotic access remains the predominant driver, an estimated 65% (∼5.7 million) of these deaths arise from barriers to access or inadequate treatment delivery[4], growing evidence indicates that even when antibiotics are appropriately used, infections frequently fail to resolve due to phenotypic modes of antibiotic survival. These include biofilm formation, tolerance, and persistence, which enable genetically susceptible bacteria to withstand transient antibiotic exposure and cause infection relapse[5]. Such phenotypic survival strategies have been repeatedly observed in clinical isolates and patient infections, contributing to recurrent and difficult-to-treat cases across multiple pathogens[6–8]. Furthermore, these non-genetic modes of survival have also been implicated as evolutionary stepping stones toward full resistance[9]. Together, these findings demonstrate that phenotypic survival represents a major, clinically relevant barrier to infection clearance, complementing and amplifying the global threat of AMR.

Despite their clinical significance, the physiological origins and dynamic transitions underlying phenotypic survival states remain poorly understood. Prior studies have examined antibiotic treatment survival in both actively dividing exponential-phase cells, where spontaneous persistence can occasionally arise[10], and in prolonged starvation or stationary-phase cells, where tolerance from dormancy is well established[11, 12]. In this study, we primarily focus on the dynamic resuscitation phase, when dormant cells awaken in the presence of antibiotics. This transition is particularly important because it combines metabolic reactivation with exposure to bactericidal drugs, creating a moment of both heightened vulnerability and heterogeneous survival strategies that can seed infection relapse. It is also highly clinically relevant, given the temporal fluctuations of nutrient availability throughout the course of an infection[13, 14], in addition to spatial heterogeneity of nutrient availability and growth-limiting host effects[15, 16], and nutrient gradients characteristic of bacterial biofilms[17].

A key feature of the resuscitation phase is the emergence of starvation-triggered persisters (Box 1), a particularly elusive and clinically important mode of nongenetic survival as described by Balaban et al.[10]. These cells survive antibiotic exposure without acquiring genetic resistance and resume growth as treatment ceases. Yet their rarity, slow growth, and transient nature make it extremely difficult to study their dynamics in bulk population assays or snapshot measurements. In addition, they are easily outcompeted by faster-dividing susceptible cells in the absence of antibiotic exposure, further obscuring their dynamics. Together, these factors have limited our understanding of how individual cells navigate resuscitation and survive antibiotic treatment, motivating the development of high-throughput single-cell approaches to observe these strategies, and predictive models to identify which survivors drive infection relapse.

To overcome these challenges, we developed MMX (Mother Machine eXtended), a next-generation microfluidic platform built on our previous high-throughput single-cell imaging platform[18], and integrated it into a complete analysis pipeline that we term Hi-DFA (High-throughput Dynamic Fate Analyser). MMX incorporates redesigned flow layouts to ensure uniform antibiotic exposure across all trenches and an innovative trench-loading architecture that enables efficient capture of individual cells (Figure S1). Isolated individual cell lineages are maintained in separate trenches, preventing fast-growing cells from outcompeting slow-resuscitating lineages, while maintaining continuous nutrient and drug exchange. The Hi-DFA pipeline couples this microfluidic setup with time-lapse microscopy and machine-learning–based image analysis, allowing automated segmentation, lineage tracking, and dynamic phenotype classification across over 100,000 individual cell lineages. Together, these capabilities provide state-of-the-art throughput and time-resolved single-cell information, which enable the quantitative mapping of the heterogeneous and reversible phenotypic transitions that shape antibiotic survival during resuscitation from dormancy. By linking each cell’s physiological history to its fate under antibiotic treatment, Hi-DFA allows a detailed and unbiased dissection of survival strategies during the clinically crucial resuscitation phase.

In this study, we focus on *β*-lactam antibiotics, including penicillins and cephalosporins, which remain the backbone of antibacterial therapy and account for more than half of global antibiotic prescriptions[19]. Because their bactericidal activity relies on active cell-wall synthesis, *β*-lactams are particularly ineffective against slow-growing or dormant cells[12, 20], a physiological state central to tolerance and persistence. To achieve a comprehensive understanding of survival strategies during resuscitation, we combined Hi-DFA with realistic *in vitro* pharmacokinetic simulations of treatment with two clinically relevant *β*-lactams, amoxicillin and cefalexin. As a model organism, we use *E. coli*, which combines well-characterised cellular physiology with direct clinical relevance: it is responsible for nearly 75% of urinary tract infections and contributes substantially to bloodstream and hospital-acquired infections worldwide[21]. By following tens of thousands of individual lineages of *E. coli* cells through the resuscitation phase under controlled antibiotic exposures, we were able to quantify how prior history of stress shapes survival strategies.

Our analysis revealed that the subpopulation typically classified as “starvation-triggered persisters” is in fact composed largely of cells that initiate regrowth upon nutrient replenishment, but demonstrate transient growth slowdown shortly thereafter. This *transient growth slowdown during resuscitation* facilitates survival during antibiotic treatment, distinguishing these cells from both *delayed-resuscitating* persisters (those with very long lag times, which we refer to as starvation-triggered persisters), and exponentially growing susceptible cells that could enter the persistence state spontaneously (which are referred to as *spontaneous persisters*)[10]. The frequency of this transiently tolerant survival phenotype, which we call “Starvation-primed transient tolerance” (see Box 1 below), scaled quantitatively with starvation duration, becoming more prevalent with increasing age of the starved culture. We thus identify, to the best of our knowledge, a previously unrecognised form of heterotolerance, present only in a subset of resuscitating cells, whose frequency is modulated by prior starvation history.

Building on these observations, and to predict which survivor classes may be the primary drivers of infection relapse, we developed a model that maps single cell outcomes to population level dynamics, capturing the diverse phenotypic trajectories encountered during antibiotic treatment and predicts population-level outcomes across varying treatment conditions. This integrative framework provides insight into the physiological determinants of survival, revealing that the presence of the rare transiently tolerant state dominates the survivor population and drives rapid infection relapse *in vitro*. Our study thus establishes both a technological and conceptual foundation for understanding and ultimately controlling phenotypic survival strategies of bacteria during clinically relevant antibiotic treatments.

### Box 1: Defining Tolerance, Heterotolerance, Persistence, and Transient Tolerance

Antibiotic tolerance describes the ability of bacterial cells to survive transient antibiotic exposure without a change in the minimum inhibitory concentration (MIC)[10]. Unlike resistance, tolerance does not alter drug susceptibility at steady state but prolongs survival during treatment. Tolerance can arise through:

- **Genetic mechanisms**, where mutations alter physiological response to antibiotics.
- **Physiological mechanisms**, where survival reflects the growth state of cells.

**Population-wide tolerance**

Beyond genetic tolerance related to specific mutations, a reversible, non-genetic reduction in antibiotic susceptibility can be observed at the bulk-population level due to physiological factors. When environmental conditions suppress growth across the entire population (e.g., nutrient deprivation, dormancy, hypoxia), most cells can become tolerant because growth-dependent antibiotic targets are inactive. Recent work[22] has demonstrated that slow or arrested growth at the population level can generate widespread tolerance without invoking phenotypic heterogeneity. In this scenario:

- The majority of cells are non-growing or slow-growing.
- Survival reflects environmental growth limitations.

**Heterotolerance**

We use the term heterotolerance to describe tolerance that emerges from phenotypic heterogeneity within a population growing under otherwise permissive conditions. In heterotolerance:

- The environment supports growth for most cells.
- A subpopulation adopts a physiological state that confers transient survival during antibiotic exposure.
- Survival arises from cell-to-cell variability rather than uniform growth suppression.

This category includes persisters and the transient tolerance phenotype described here.

A. **Starvation-triggered persistence:** Persisters are a subpopulation of cells that enter a prolonged non-growing or extremely slow-growing state prior to antibiotic exposure. Persisters are typically associated with long lag times following starvation or stress. Starvation-triggered persisters refer to the subpopulation of cells that demonstrate extended lag and low growth rates from the start of resuscitation, and delayed regrowth after antibiotic removal[10].
B. **Starvation-primed transient tolerance:** We define transient tolerance as a distinct heterotolerant state arising during resuscitation from starvation. Starvation-primed transient-tolerant cells refer to the subpopulation of cells that initially demonstrate growth rates comparable to the susceptible (or untreated) population at the start of resuscitation, but subsequently experience a transient growth slowdown that facilitates survival during *β*-lactam antibiotic treatment. Notably, these transient growth slowdown events occur even in the absence of treatment, and their prevalence is scaled by starvation duration. Unlike persisters, transiently tolerant cells:
  - Are not deeply dormant.
  - Do not exhibit prolonged lag.
  - Rapidly rebound after treatment.

## 2 Results

### 2.1 Hi-DFA: A High-Throughput Platform For Detecting and Tracking Rare Antibiotic Tolerant and Persistent Bacterial Cells

Probing bacterial dynamics of phenotypic survival states in response to antibiotic treatment requires characterisation of rare and transient events, such as antibiotic persistence. Identifying and accurately quantifying such rare events at the single-cell level hence requires very high throughput. Therefore, we developed MMX, an iteration on the mother machine microfluidic device[23] that allows for time-resolved imaging of up to 115,200 individual bacterial cell lineages in parallel, across up to 6 different conditions. The density of the device allows us to image at time resolutions as high as 1–2 minutes and low spatial resolutions (at 30× magnification and 0.75 NA, Figure 1). This level of throughput was achieved by compact trench organisation and redesigning the flow layouts into an extensive “three-lane-snake” design (Figure 1a,b) that maximises the number of trenches in each lane whilst ensuring uniform conditions throughout the entire lane. This design of three individual lanes arranged as a serpentine was chosen to improve upon previous devices with a larger number of serpentined lanes, which results in a depleting antibiotic and nutrient concentration along the lane, unsuitable for concentration-sensitive conditions such as treating with low concentrations of antibiotics[18]. Multiple lanes per device also provide the advantage of testing up to 6 different conditions in parallel (Figure 1a), providing increased consistency between treatment conditions. The MMX devices also introduce a novel trench-loading architecture by incorporating structured large surface-area void spaces above each cell trench (Figure S1). This void space generates suction driven by evaporation from the partially permeable PDMS[24] backports (Figure 1c,d). Evaporation through PDMS also occurs in conventional mother machine devices. In MMX, the thin PDMS membrane above the backport concentrates this evaporative flux into a structured channel directly above each trench, creating a directed flow that actively draws cells into the trench. This geometry significantly improves the efficiency of trench loading and increases cell retention duration (Figure S2, Video 1), providing the highest possible throughput and reducing the wait time during experimental setup.

**Figure 1:**
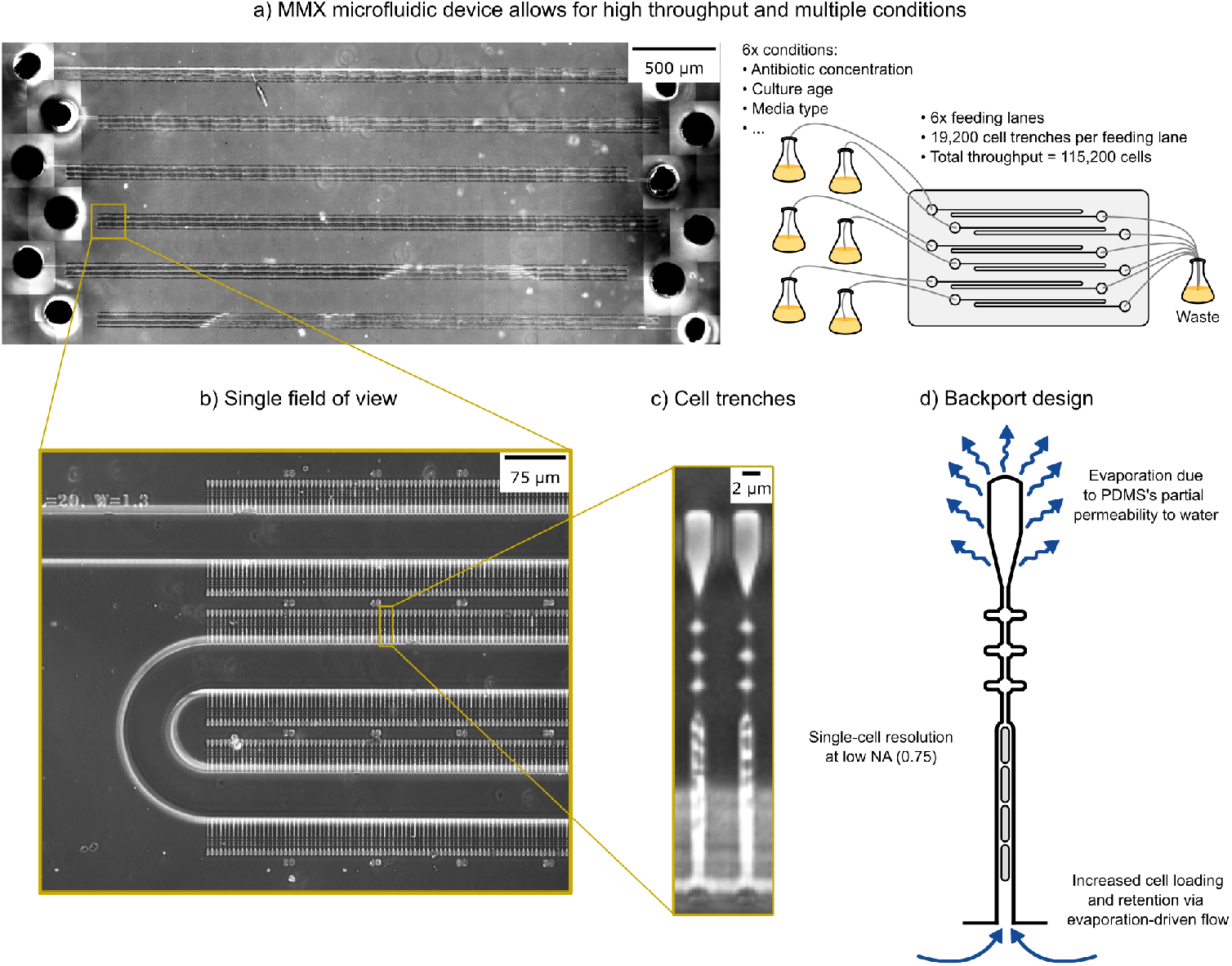
MMX: A high-throughput microfluidic platform for detecting and tracking rare antibiotic tolerant and persistent bacterial cells. **a)** Scan of the entire MMX device with phase contrast imaging. The MMX device allows for high throughput while also maintaining the ability to test multiple (six) conditions simultaneously, with a throughput of 19,200 cells each (for a total of 115,200 cells). This throughput is sufficient to derive accurate measurements of very rare events and phenotypes. **b)** A single field of view (FOV) of the MMX device shows the nutrient feeding lane’s geometry. **c**,**d)** Individual cell trenches are modified with a backport design which promotes evaporative flow from the closed end. This boosts cell-loading speeds and cell retention which are critical for high throughput measurements.

To analyse the extensive single-cell antibiotic treatment response data generated from experiments utilising the MMX devices, we developed an image analysis pipeline, subsequently referred to as Hi-DFA (High-throughput Dynamic Fate Analyser), combining trench identification and extraction with machine-learning-based cell segmentation, cell feature extraction, and individual cell fate characterisation (Figure 2, Supplementary Note 1). This pipeline involves first training a machine-learning model for accurate segmentation of bacterial cells from the microscopy images. This was achieved using the Omnipose architecture[25], initially trained on synthetic microscopy images generated by SyMBac[26], then fine-tuned on curated well-segmented data from applying the SyMBac trained model to experimental microscopy images from MMX devices (Figure 2a). Following fine-tuning of the segmentation model, individual trenches from the microscopy images were identified (Figure 2b), then cropped and assembled into compact arrays for subsequent processing (Figure 2c,d). Note that the typical dataset contains hundreds of fields of view with each field of view containing hundreds of trenches. Utilising Zarr[27] for on-disk storage and Dask[28] for processing facilitates efficient analysis of these large files. This allows Hi-DFA to run on readily available consumer hardware where memory is often constrained. To promote accessibility and reproducibility, the complete Hi-DFA analysis pipeline has been published on GitHub[29].

**Figure 2:**
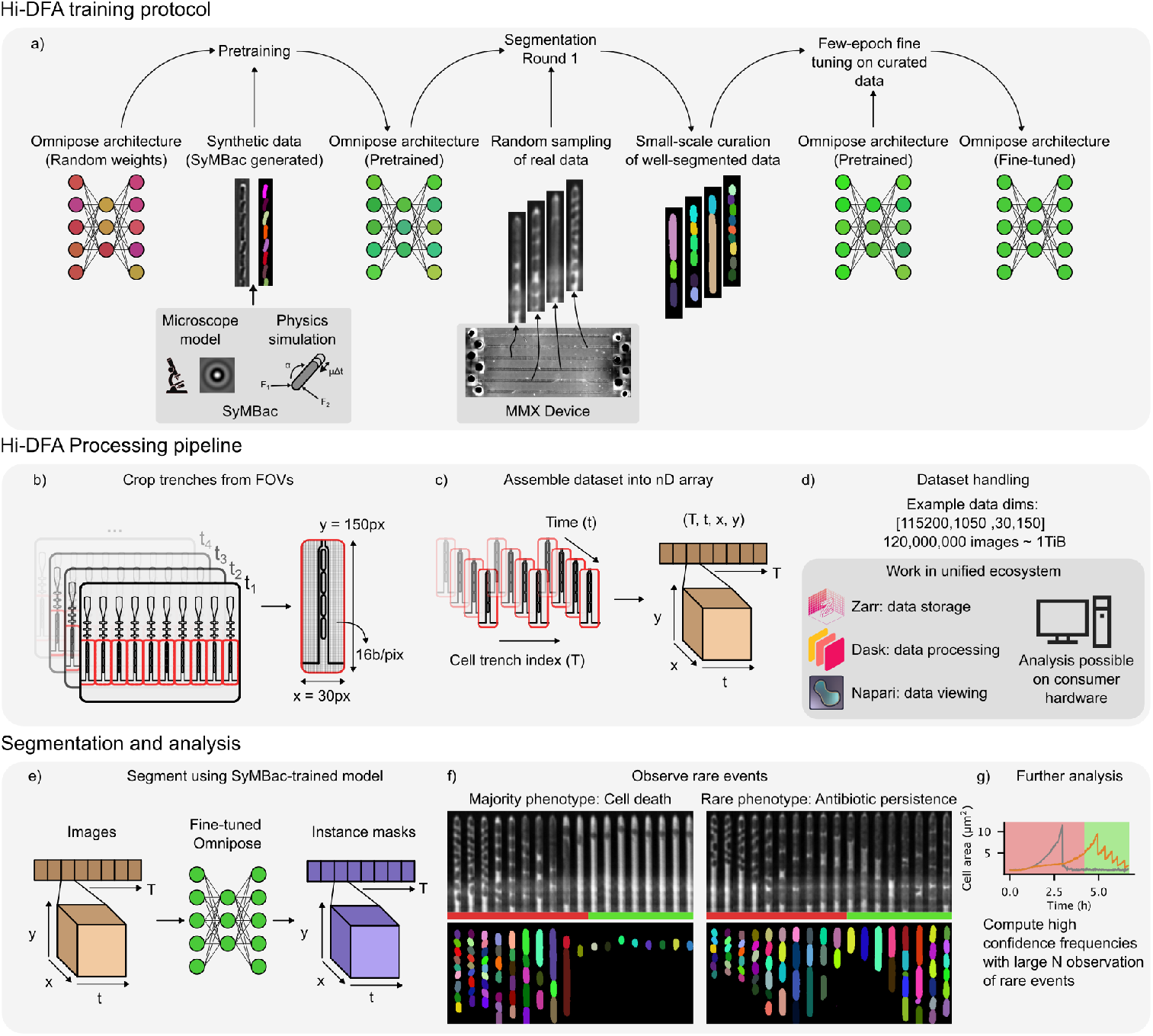
Automated data processing and machine-learning pipeline for dynamic single-cell fate analysis in Hi-DFA. **a)** The Hi-DFA training protocol involved pre-training the Omnipose model on large amounts of synthetic training data[26], which was followed by an initial round of segmentation of a small sample of experimental data. The segmentation results were then manually curated (selected/rejected) for fine tuning the model. This resulted in a model highly capable of segmenting low-resolution images with a high degree of precision. **b)** The Hi-DFA processing pipeline involves extracting raw data from FOVs into individual crops of single MMX cell trenches. **c)** These cell-trenches crops are then stacked by index and time, and stored in an nD array. **d)** By working entirely in the mature Python data-science ecosystem, one can manipulate datasets as large as 1 TiB on consumer hardware. This is facilitated by lazy loading of nD data stored on disk in Zarr[27] arrays (a format to store multidimensional arrays which is used widely in the sciences), processing using Dask[28], which implements automatic implementation and efficient lazy execution of computation graphs, and Napari[30], which can view these results, also lazily. **e)** The entire MMX dataset can be segmented using the fine-tuned model, with the masks stored as Zarr on disk, also allowing for convenient and efficient processing. **f)** Analysis of the resulting images and masks enables observation of rare events. For example, most cells die upon exposure to antibiotics (left kymograph), but a rare fraction are able to tolerate high concentrations for several hours and resume growth after the transient exposure (right kymograph). **g)** Growth traces corresponding to the area of the mask of the mother cell in the examples in f). Cell lysis (grey) vs survival (green) under antibiotic treatment (red shaded area). These time series data can then be grouped and compared across different treatment conditions to yield biological insights.

To extract and track the phenotypes of individual cell lineages, cropped images corresponding to individual trenches were input to the fine-tuned machine-learning segmentation model to generate cell masks for each time point. Masks from consecutive frames were subsequently registered and linked to reconstruct single-cell trajectories, from which temporal features such as growth rate, division events, and lysis events were extracted for dynamic fate analysis (Figure 2e). The ultimate fate of the mother cell following antibiotic treatment can be determined from the mask features, providing dynamics of cell death that occurs for the majority of the population, in addition to identifying rare events such as persistence in a subset of the population (Figure 2f). Hi-DFA analysis of different treatment conditions within or between experiments therefore allowed us to (i) distinguish different types of antibiotic treatment survivor phenotypes, (ii) quantify the frequencies of different classes of survivors, to determine which are the most prevalent, and (iii) to determine how the frequencies of these survivor classes are affected by the environment and different antibiotic treatment conditions.

### 2.2 Hi-DFA Reveals a Novel Heterotolerance Phenotype (Transient Tolerance) Unlocked by Prior Stress

Using the Hi-DFA pipeline, we studied the dynamic cell fates of resuscitating *E. coli* under *β*-lactam antibiotic treatment. We primarily focused on treatments occurring in two growth regimes: resuscitation from stationary phase and exponential growth (Figure 3a). Antibiotic treatment during exponential growth has frequently been studied in previous work on antibiotic persistence[10, 20] and provides an insight into mechanisms that occur when cells are experiencing minimal amounts of stress and reduced “memory” of that stress from prior starvation. Treatment during the resuscitation period provides important insights into mechanisms that may be affected by metabolic reactivation or stresses experienced during starvation. Unlike in bulk population experiments, single-cell-level analysis of antibiotic treatment during the resuscitation regime allows us to differentiate survival phenotypes that are caused by starvation from survival phenotypes that are caused by the antibiotic, facilitating improved understanding of survival modes during this clinically crucial phase. Antibiotic exposure of 50 µg/mL ampicillin (∼3× MIC) for four hours demonstrated that treatment during the resuscitation period resulted in greatly increased survival frequency relative to treatment during the exponential growth phase (Figure 3b,c), consistent with previous studies[10, 20].

**Figure 3:**
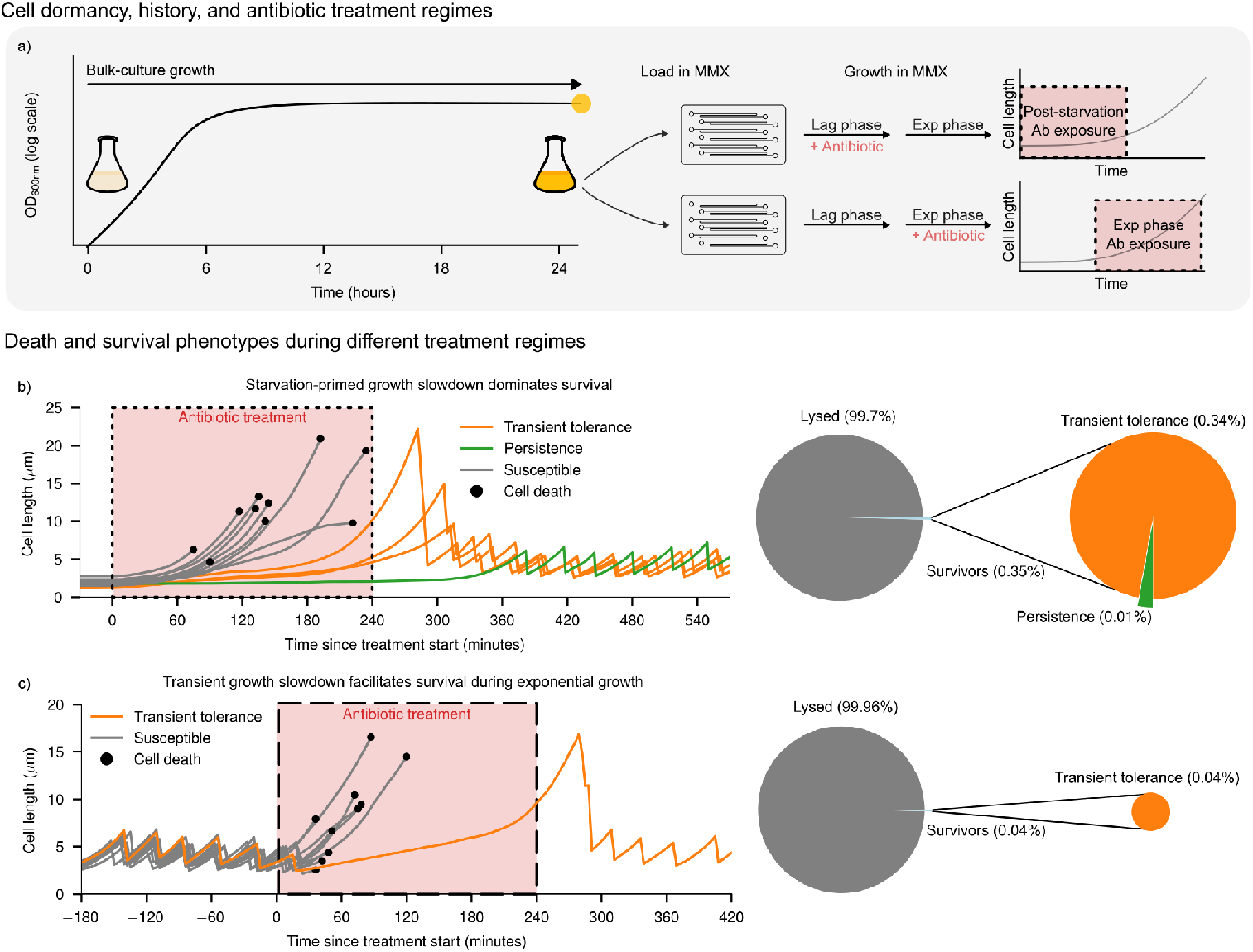
A high-throughput platform reveals a new antibiotic treatment survivor phenotype primed by starvation. **a)** Schematic of bacterial culture cell number (OD_600_) over time for a bulk culture of bacteria entering and remaining in stationary phase for an extended period, before loading into the MMX device and cell awakening following exposure to fresh media. The regions highlighted in red indicate antibiotic treatment occurring at either of the two major growth regimes explored in this paper—immediately during awakening, during post-starvation bacterial resuscitation, or later on, during steady-state exponential growth. **b)** Left: Representative single-cell length traces of resuscitating cells waking up from stationary phase in the presence of an antibiotic. The cells either lyse during antibiotic treatment (grey traces), survive due to starvation-primed growth slowdown after engaging in normal population-level resuscitation kinetics (orange traces), or survive due to starvation-triggered persistence (green trace). The red shaded region indicates the timing of 4 hours of treatment with 50 µg/mL ampicillin. Right: Distribution of cell fates within the population following antibiotic treatment. Number of cells = 22,219. **c)** Left: Representative single-cell length traces of cells in the exponential growth phase that either lyse during antibiotic treatment (grey traces) or survive due to transient growth slowdown (orange trace). The red shaded region indicates the timing of 4 hours treatment with 50 µg/mL ampicillin. Right: Distribution of cell fates within the population following antibiotic treatment. Number of exponentially growing cells = 37,167.

During antibiotic treatment of a resuscitating 24-hour-old culture, the vast majority of cells (99.7%) exited dormancy, resumed growth, and eventually lysed (grey, Figure 3b). Among the surviving fraction (0.35%), a small subset (0.01%) consisted of cells that remained semi-dormant throughout the treatment period, demonstrating little or no increase in size (green, Figure 3b). Their survival can be attributed to extended and heterogeneous lag times prior to resuming exponential growth, a behavior consistent with classical starvation-triggered persisters[10]. In contrast, the dominant class of survivors (0.34% of the entire population, that is, >97% of survivors) displayed a distinct phenotype: these cells initially grew at rates comparable to the susceptible population, but shortly after resuscitation in the presence of the antibiotic, they demonstrated a transient decrease in growth rate, facilitating treatment survival (orange, Figure 3b). Notably, these cells usually did not demonstrate complete growth arrest; instead, they maintained a transiently reduced but non-zero growth rate for a variable duration, followed by a subsequent rapid increase in growth rate (Video 2). These growth dynamics clearly differentiate this class from classical starvation-triggered persisters, which maintain very low growth rates from the start of treatment (Figure 3b). In the following sections, we characterize this transiently slow-growing subpopulation in detail and show that its emergence depends on prior starvation history and on the cellular response to antibiotic exposure. We therefore designate these cells as *starvation-primed transiently tolerant* cells.

In contrast, treatment during the exponential growth phase resulted in near-complete killing of the population. The small number of cells that survived (0.04%) demonstrated a transient decrease in growth rate during antibiotic treatment, facilitating their survival (Figure 3c, Figure S3). Notably, in all cases where we studied antibiotic exposure of already exponentially growing cells, this decrease in growth rate initiated spontaneously prior to treatment and continued to decrease further following treatment start (Figure S3); further characterisation of this spontaneous transient tolerance phenotype during exponential growth is required, but beyond the scope of this study which focuses primarily on the exit from stationary phase. Regardless, this treatment survival during exponential growth (owing at least partially to rare spontaneous growth slowdown) appears consistent with the population-level classification of *spontaneous persisters*[10]. However, we note that these cells do not exhibit complete growth arrest (Figure S3); rather, they continue to grow at a reduced rate. The very low frequency of this survivor class during the exponential growth phase suggests that the growth slowdown of transient tolerance is primarily programmed during starvation and gradually erased during subsequent growth (Figure 3c). Survival during resuscitation occurred at ∼0.35%, compared to ∼0.04% in exponential growth—an approximately tenfold difference. This substantial separation demonstrates that transient growth slowdown is markedly enriched during resuscitation from starvation, consistent with the interpretation that transient tolerance is programmed by prior stress rather than arising spontaneously at appreciable frequency.

### 2.3 Starvation-Primed Transient Tolerance is Distinct from Starvation-Triggered Persistence

The susceptible class of cells starts and maintains exponential growth in cell size soon after resuscitation until eventually lysing due to the presence of the antibiotic (Figure 4a). Notably, the transiently tolerant class initially starts growing at a comparable rate to the susceptible population, before experiencing a transient slowdown shortly after the start of antibiotic exposure (30–120 min), followed by resuming the rapid increase in size that facilitates cell division events following the end of treatment (Figure 4a,b). The similar growth rates between the susceptible cells and the starvation-primed surviving subpopulation of bacteria at the start of resuscitation demonstrate that these survivors appear to emerge from within the susceptible population (Figure S4). Notably, transient growth slowdown events were also observed in resuscitating cells in the absence of treatment (Figure S5), indicating that this behaviour is encoded by stress history, not solely induced by antibiotic exposure, though the onset timing may be modulated by treatment. Similarly, their regrowth is independent of antibiotic removal, and widely distributed. In contrast, starvation-triggered persisters maintain a low (but usually non-zero) growth rate from the start of resuscitation (Figure 4a,b). The frequency of this class also does not seem to be affected by antibiotic exposure (Figure S6).

**Figure 4:**
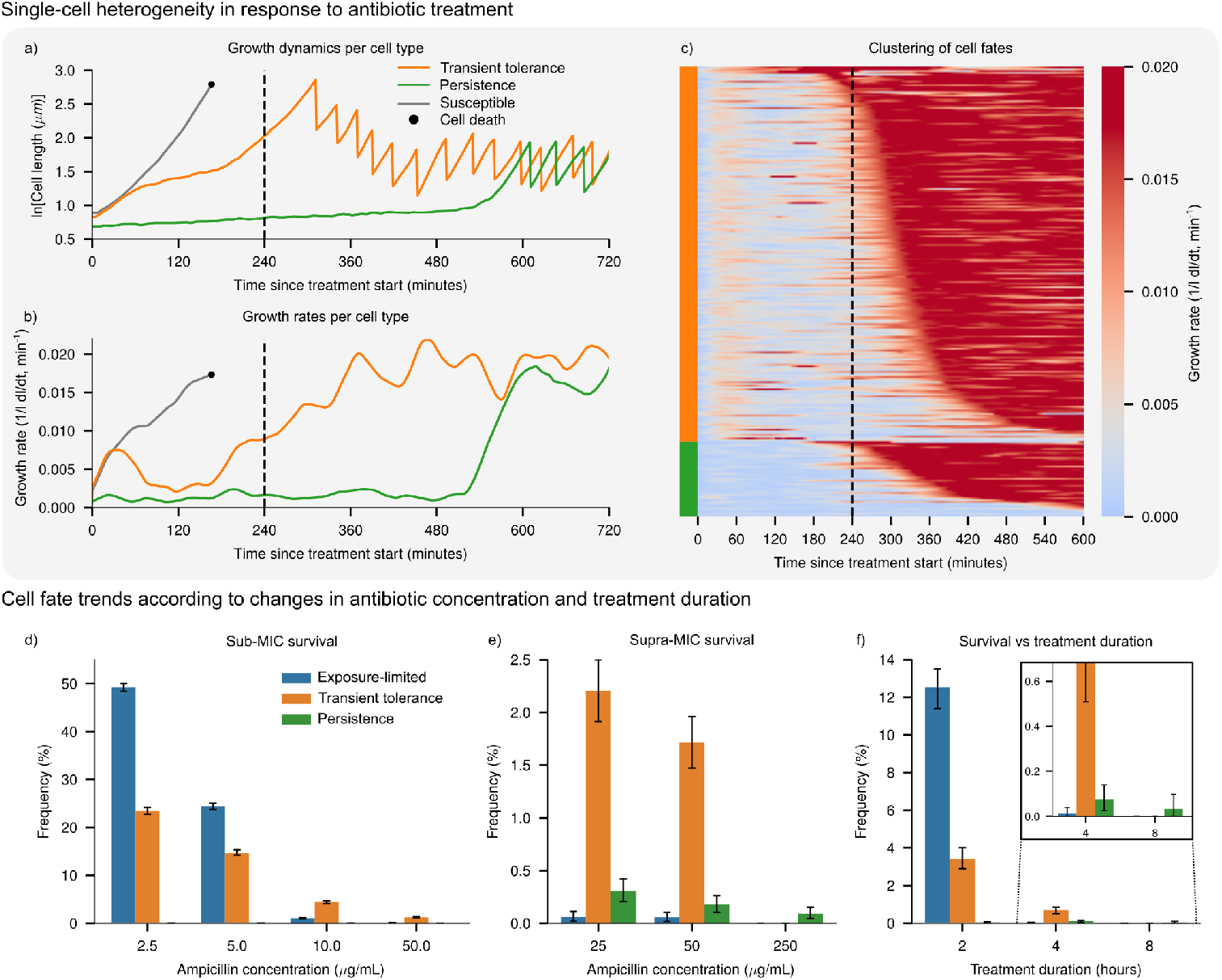
Transient tolerance is distinct from classical persistence. **a)** Representative single-cell length traces (log scale) of three classes of cells resuscitating from stationary phase in the presence of an antibiotic. The vertical black dashed line indicates the end of 4 hours treatment with 50 µg/mL ampicillin. **b)** Representative single-cell growth rate traces corresponding to the derivative of the cell length traces in (**a**). **c)** Combined single-cell growth rate traces of 382 surviving cells from a single experiment. Traces were clustered into 2 clusters from the growth rate trace at the start of treatment, then sorted within each cluster by onset time of high growth rate increase. **d)** Frequencies of the three classes of surviving cells following 4 hours treatment with the specified sub-MIC concentration of ampicillin (MIC = 16 µg/mL). **e)** Frequencies of the three classes of surviving cells following 4 hours treatment with the specified supra-MIC concentration of ampicillin. **f)** Frequencies of the three classes of surviving cells following treatment with 50 µg/mL ampicillin for the specified duration. Error bars indicate the upper and lower 95% confidence intervals from bootstrapping with *n* = 10,000.

To determine whether transiently tolerant cells represent a distinct physiological state, we compared the single-cell growth dynamics of all survivors. Clustering analysis of over 300 survivor growth-rate trajectories revealed two clearly separable groups: one displaying an initial rise followed by a transient decline in growth during treatment, and another remaining largely growth-arrested until much later recovery (Figure 4c). There is substantial heterogeneity in the time at which transiently tolerant survivors resume normal growth and division, with Figure 4c showing this class of cells ranked by the time at which the mean balanced growth is resumed. Conversely, we find a relatively lower heterogeneity in the actual dynamics of the initial slow-down in survivors, with most cells behaving somewhat similarly, exiting stationary phase at rates commensurate with the rest of the population, and then slowing down their growth to a minimum at approximately 120 minutes post-exposure. Furthermore, the heterogeneity in the resumption to full growth appears to be uncorrelated with the treatment duration. This is evident from the wide variation of their growth resumption dynamics shown in Figure 4c, but also with the fact that increasing treatment duration (Figure 4f) leads to increased killing of this class.

Next, we investigated how the frequency of each survivor class was affected by the antibiotic concentration and treatment duration. At concentrations of ampicillin far below the MIC, the majority of surviving cells did not exhibit notable growth slowdown (Figure 4d). These survivors fall within the susceptible population and likely reflect intrinsic heterogeneity in resuscitation timing or early growth rate. Cells with slightly longer lag durations or lower initial growth rates may remain below the threshold for effective growth-dependent killing during brief or low-dose treatment, not through a programmed tolerance mechanism, but because antibiotic exposure is insufficient relative to their growth trajectory (Figure S4). We therefore refer to these as **exposure-limited survivors** (defined in Box 2). However, as the antibiotic concentration increased, the frequency of exposure-limited survivors dropped rapidly and transiently tolerant survivors became the dominant class of survivors (Figure 4d,e).

#### Box 2: Exposure-limited survivors

Cells that survive antibiotic treatment due to limited exposure duration despite exhibiting the default susceptible phenotype. These cells arise from natural variability in lag time within the susceptible population: a subset of cells exit lag more slowly and therefore do not reach the growth-dependent threshold required for antibiotic-mediated killing before treatment ends.

As the treatment concentration of ampicillin approaches the MIC, the total surviving fraction of cells following 4 hours of treatment reaches ∼2%. At this point, transient tolerance, programmed by past stress from starvation, becomes the dominant driver of survival amongst the remaining cells. However, as the concentration increased further beyond the MIC, the frequency of these survivors decreased until at concentrations well above the MIC (>15× MIC) complete killing of the transiently tolerant class was achieved and starvation-triggered persisters comprised the dominant class of survivors (Figure 4e). The survival of the starvation-triggered persisters at these high antibiotic concentrations has previously been attributed to the killing rate of *β*-lactam antibiotics being proportional to growth rate[12, 20], and we emphasise that at the single-cell level, this is fundamentally equivalent to survival-by-lag: cells that delay growth initiation avoid susceptibility to growth-dependent killing until antibiotic levels decline. A similar trend of survivor class frequencies was identified for varying antibiotic treatment duration.

Short durations (2 hours) of 50 µg/mL ampicillin (∼3× MIC) resulted in exposure-limited survivors comprising the majority of survivors (Figure 4f). As treatment duration increased, transiently tolerant survivors became the most prevalent class of survivors, until at long treatment durations (8 hours), starvation-triggered persisters were the dominant class of survivors (Figure 4f). Thus, when antibiotic exposure is sufficiently long or high concentration, survival becomes primarily determined by lag duration rather than transient growth modulation during resuscitation. These results demonstrate that, despite both being triggered by history of starvation, starvation-primed transient tolerance and classical starvation-triggered persistence are distinct classes of survival that differ both in growth dynamics during antibiotic treatment, and in prevalence and susceptibility to concentration and duration of antibiotic treatments.

Notably, transiently tolerant cells remain actively growing during antibiotic treatment, albeit at reduced rates. Many of these cells elongate substantially under *β*-lactam exposure, consistent with filamentation. At the end of treatment, a subset of transiently tolerant survivors are markedly elongated. Upon antibiotic removal, these filamented cells undergo rapid septation, generating multiple progeny from a single surviving lineage (orange line, Figure 4a). In contrast, classical persisters typically remain growth-arrested during treatment and resume division only after an extended lag. As a result, each transiently tolerant survivor can contribute more than one descendant cell immediately following washout, effectively increasing the reproductive output per surviving lineage and accelerating early population rebound. This morphological amplification mechanism, detailed in Figure S7, is fully concordant with the single-cell length and division trajectories captured by Hi-DFA, which show a transient growth slowdown followed by a rapid size increase and subsequent division bursts after washout.

### 2.4 Starvation Duration Quantitatively Modulates Transient Tolerance Frequency

Since these transiently tolerant cells were frequent in starved populations but virtually absent in exponentially growing populations, we hypothesized that their emergence is primed by the history of starvation. To investigate the effect of starvation duration on the frequencies of different survivor classes following antibiotic treatment, we carried out ampicillin treatment of bacterial cells that were either actively growing in exponential phase, or treatment of cells resuscitated from stationary phase cultures of different ages—24, 48, or 72 hours old (Figure 5a).

**Figure 5:**
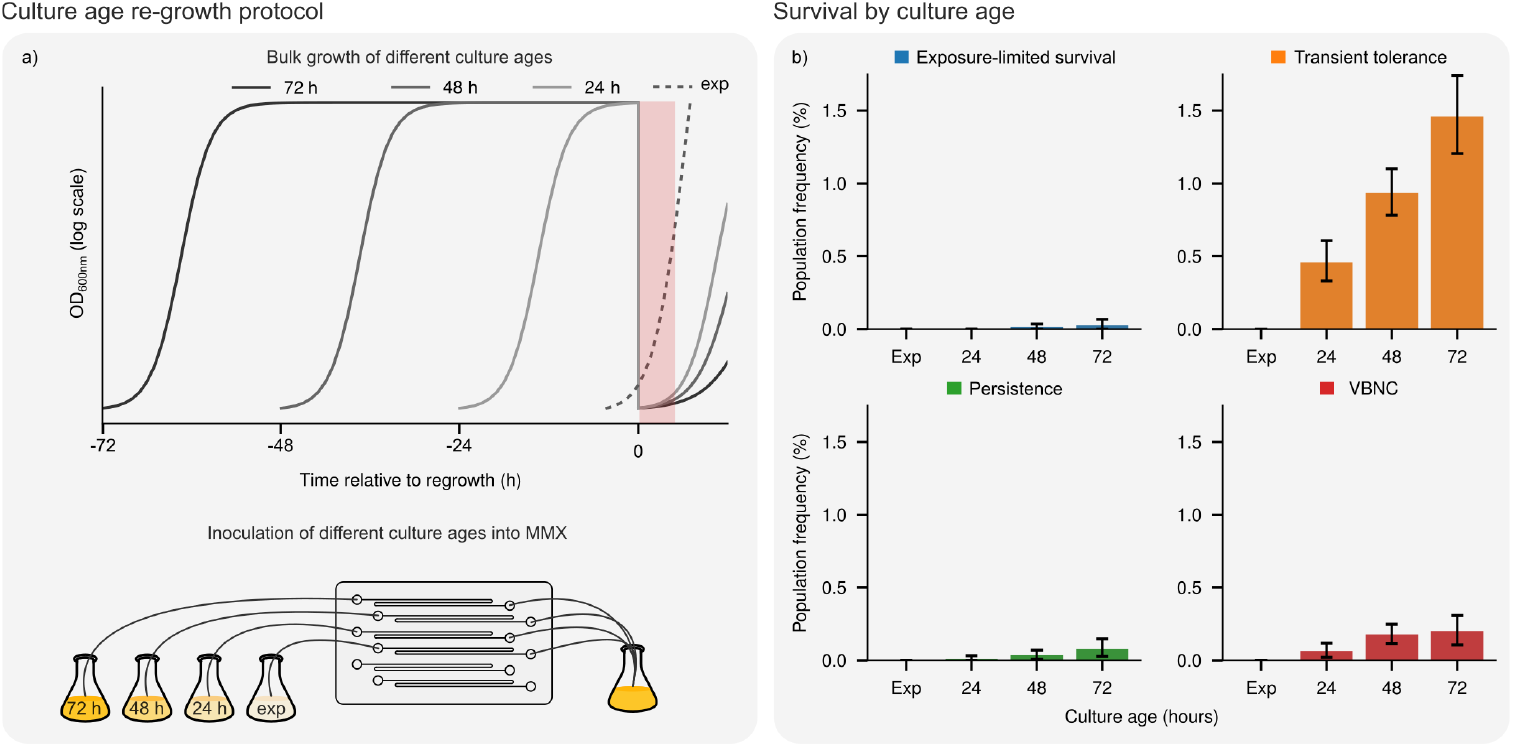
Starvation duration quantitatively modulates transient tolerance frequency and cell fate during resuscitation. **a)** Schematic of bacterial culture inoculation timing prior to antibiotic treatment. The red shaded region indicates 4 hours treatment with 50 µg/mL ampicillin. **b)** Frequencies of four classes of surviving cells following 4 hours treatment with 50 µg/mL ampicillin. The culture age on the *x*-axis refers to the age of the bacterial cultures at the time of antibiotic treatment. The number of cells for each culture age are (Exponential phase: 6,506, 24 hr old: 9,387, 48 hr old: 14,077, 72 hr old: 7,471). Error bars indicate the upper and lower 95% confidence intervals from bootstrapping with *n* = 10,000.

Treatment of these differently aged cultures identified that increased starvation duration prior to resuscitation tended to increase the frequency of all survivor classes (Figure 5b). The increased frequency of starvation-triggered persisters and viable but nonculturable (VBNC) bacteria[31] (cells that remain viable but fail to resuscitate during the observation period) with increasing starvation duration is consistent with previous reports[20, 32, 33]. Their extended lag times and slow or impaired resuscitation are likely linked to factors such as cellular ATP depletion during starvation[34].

However, as starvation duration increased, the increase in survivor frequency was most pronounced for the transiently tolerant class of survivors, which comprised the majority of antibiotic treatment survivors, particularly in the older cultures (Figure 5b). This suggests that starvation experienced during stationary phase leaves a measurable duration-dependent effect on the dynamics of resuscitating cells. This prior physiological history quantitatively determines the likelihood of an individual cell undergoing transient growth slowdown and thereby determines its fate upon antibiotic exposure during the resuscitation period.

### 2.5 Clinically Relevant Pharmacokinetic Profiles Select for Transient Tolerance

After demonstrating that the prevalence of different survivor classes was significantly affected by antibiotic treatment concentration and duration, we aimed to identify the prevalence of the different survivor classes under clinically relevant antibiotic treatment pharmacokinetics. While fixed-concentration experiments are useful for isolating variables, they do not represent the dynamic concentration changes that occur in a patient’s body following dosing. These clinical pharmacokinetic (PK) profiles, characterized by a peak concentration (*C*_max_ at time *T*_max_) followed by an elimination phase (*t*_1/2_), expose bacteria to ever changing stress.

Understanding how survivor phenotypes emerge under these realistic, time-varying antibiotic pressures is therefore essential for translating *in vitro* findings to clinical scenarios. To achieve this, we built a multi-pump system to precisely control the concentration of antibiotic flowing into the MMX device over time (Figure 6a, details in Supplementary Note 2). This system allowed us to mimic the typical clinical pharmacokinetic profiles for single doses of common antibiotics (Figure 6b), providing increased clinical relevance over fixed antibiotic concentrations over a period of time. From this, we identified that treating bacteria during the resuscitation period with antibiotic concentrations mimicking the pharmacokinetic profiles of two commonly prescribed[35, 36] *β*-lactam antibiotics, amoxicillin[37, 38] (Figure 6c) or cefalexin[39] (Figure 6e), demonstrated that single doses of these common antibiotics are ineffective at complete killing, and that the survivors were dominated by the transiently tolerant class (Figure 6d,f).

**Figure 6:**
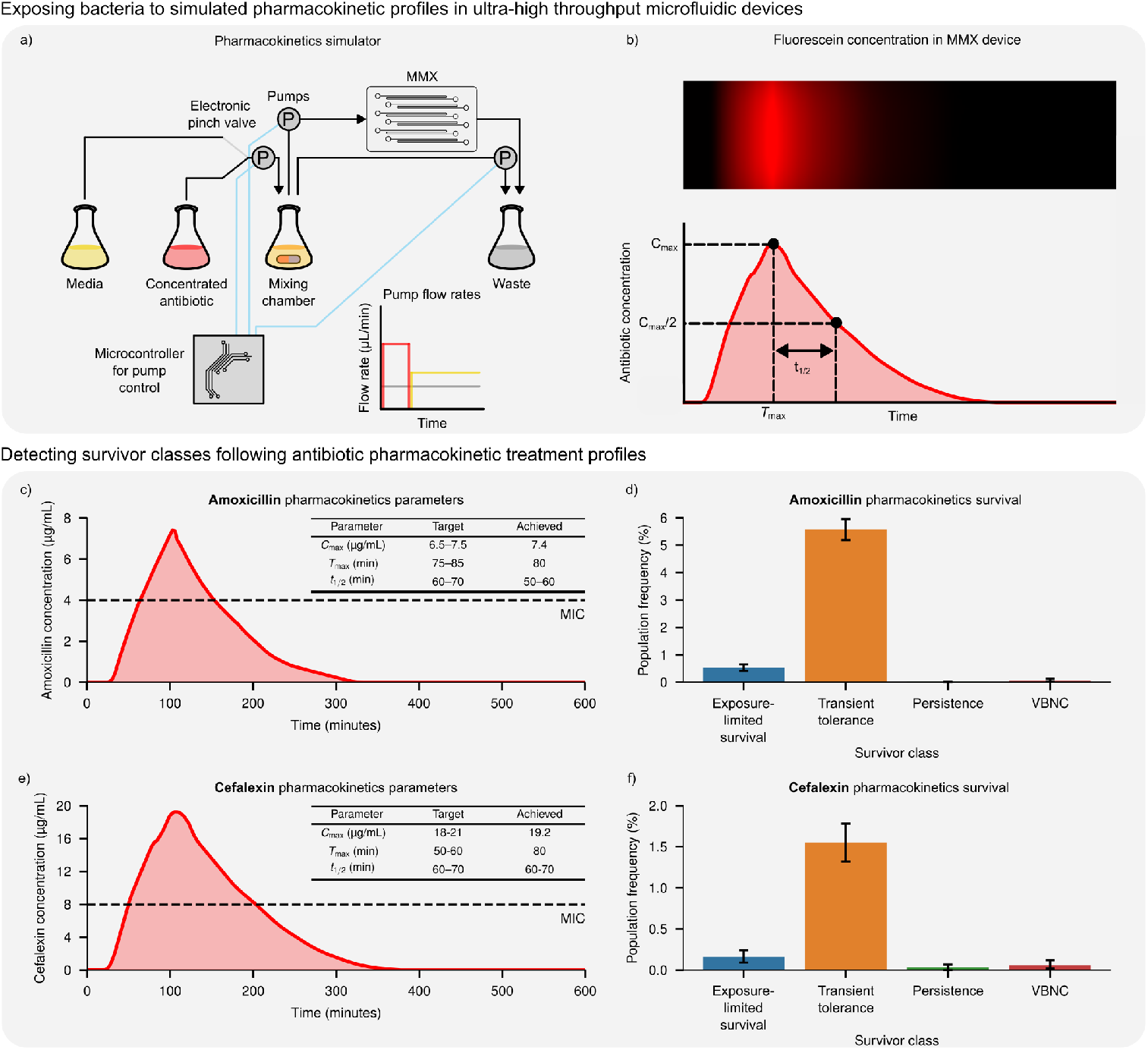
Clinically relevant pharmacokinetic profiles select for transiently tolerant cells. **a)** Schematic depicting the multi-pump system for mimicking clinical antibiotic pharmacokinetics within the MMX device. **b)** Top: Kymograph of the mean green fluorescence intensity within a feeding lane of the MMX device for the first 600 minutes of amoxicillin pharmacokinetics. Bottom: Example pharmacokinetic profile after a single antibiotic dose. *C*_max_ refers to the peak antibiotic concentration reached in the target compartment (e.g. blood plasma). *T*_max_ refers to the time taken from the start of treatment to reach *C*_max_. *t*_1/2_ refers to the approximate time taken for the antibiotic concentration to decrease by 50%. **c)** Experimentally achieved amoxicillin concentration profile for mimicking clinical amoxicillin pharmacokinetics. The horizontal dashed black line indicates the MIC of 4 µg/mL. **d)** Frequencies of the four classes of surviving cells after amoxicillin PK antibiotic treatment from c). Number of cells = 13,619. **e)** Experimentally achieved cefalexin concentration profile for mimicking clinical cefalexin pharmacokinetics. The horizontal dashed black line indicates the MIC of 8 µg/mL. **f)** Frequencies of the four classes of surviving cells following the cefalexin PK antibiotic treatment from e). Number of cells = 10,030. Error bars indicate the upper and lower 95% confidence intervals from bootstrapping with *n* = 10,000.

The high frequency of transiently tolerant survivors, over 50-fold greater than the frequency of classical starvation-triggered persisters, following treatment with these clinically relevant pharmacokinetic profiles of common antibiotics suggests that transient tolerance may be a major driver of *β*-lactam antibiotic treatment failure and infection recurrence. Notably, when treated with a different *β*-lactam antibiotic, ceftriaxone, antibiotic concentrations 10× the MIC resulted in significant prevalence of transiently tolerant survivors, but clinically relevant concentrations of ceftriaxone resulted in complete killing (Figure S8), likely owing to the clinical peak serum concentration of ceftriaxone being >10,000× the MIC of the used *E. coli* strain[40, 41] (Table S1). Hence, complete killing following *β*-lactam antibiotic treatment can be achieved through antibiotic choice, and perhaps also through combining treatment with appropriate adjuvants or by modifying the antibiotic’s pharmacokinetic profile, described below in the discussion.

Antibiotics such as amoxicillin and cefalexin are typically prescribed for multiple doses per day[38, 42, 43], hence survival frequencies from a single dose of treatment are unlikely to be representative of the clinical outcome. However, given the high prevalence of the transiently tolerant survivor class and the fact that they are actively growing at the end of antibiotic treatment, these cells could likely significantly increase in number in the time between treatment doses, hence limiting the efficacy of multi-dose treatments.

### 2.6 Transient Tolerance Dominates Rapid Population Rebound and Drives Treatment Failure

Our experimental data, particularly from clinically relevant pharmacokinetic profiles, demonstrate that transiently tolerant survivors frequently dominate the surviving population after a single antibiotic dose. We also note that at longer treatment times or higher antibiotic concentrations, persisters become the dominant surviving subpopulation (Figure 4f). However, survivor frequencies alone do not reveal which cell class is ultimately responsible for population regrowth, since the relapse potential of a class is also determined by its ability to resume and maintain fast growth post-treatment. Therefore, a critical unanswered question is, in treatment conditions not analysed in this study, is treatment failure primarily driven by this larger population of fast-resuscitating, transiently tolerant cells or by the smaller, slow-growing fraction of starvation-triggered persisters?

Answering this question requires a framework that connects the growth physiology of individual cells to their probability of surviving treatment. Because *β*-lactam killing is mechanistically coupled to active cell-wall synthesis, the rate at which a cell accumulates lethal damage is approximately proportional to its instantaneous growth rate. The total damage sustained over the treatment window is therefore proportional to the time-averaged growth rate during exposure, *λ*_avg_, and a cell dies when this cumulative damage exceeds a lethal threshold[44]. This defines a critical *λ*_avg_: cells whose time-averaged growth rate falls below this threshold survive, while those above it accumulate fatal damage (Figure 7a,b). Importantly, this framing is consistent with previous observations that total added cell length during *β*-lactam exposure predicts single-cell survival probability[44].

**Figure 7:**
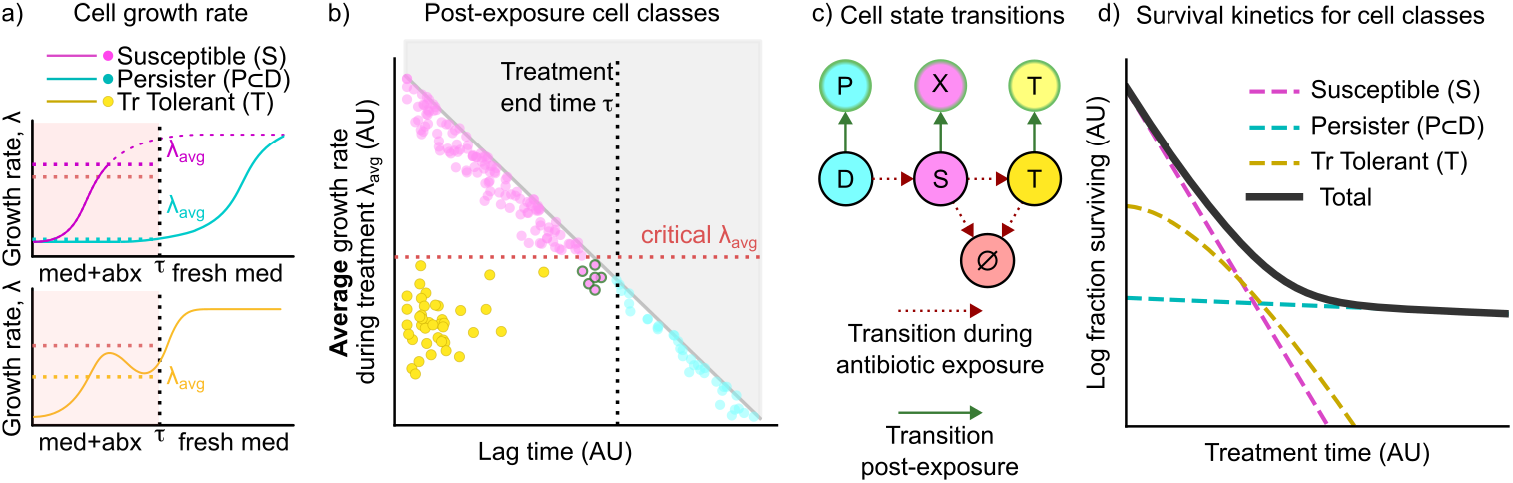
Transient tolerance produces a distinct intermediate phase in population kill curves. **a)** Schematic growth rate (*λ*) trajectories for cells exiting the stationary phase upon the addition of fresh media and antibiotics. Shown are susceptible (*S*, magenta), transiently tolerant (*T*, yellow), and persister (*P* ⊂ *D*, cyan) cell classes during and after antibiotic treatment. **b)** Classification of post-exposure cell fates in *λ*_avg_–lag time space (schematic representation). Cells with lag time *L* > *τ* (right of vertical dashed line) survive as persisters (*P* ⊂ *D*). Among resuscitated cells (*L* ≤ *τ*), those above the critical *λ*_avg_ (horizontal red dashed line) accumulate lethal damage (magenta), while those below it survive as transiently tolerant cells (*T*, yellow). **c)** Cell state transition diagram for both antibiotic exposure (red dashed arrows) and post-exposure (green solid arrows). **d)** Schematic survival kinetics illustrating how the three cell classes contribute to the overall kill curve.

During resuscitation from the stationary phase, cells exhibit heterogeneous lag times (*L*) before initiating growth. Once growth resumes, the instantaneous growth rate increases gradually toward the maximum, so that cells which exit dormancy earlier spend more time growing rapidly and thus achieve a higher *λ*_avg_ than those that exit later (Figure 7a). Crucially, this relationship is constrained by recovery physiology: cells cannot exit dormancy late and immediately achieve a maximal growth rate, since growth acceleration requires prior metabolic recovery (grey inaccessible region, Figure 7b). Without any additional survival mechanism, this constraint places all resuscitating cells within a boundary in the *λ*_avg_–lag time space (Figure 7b), where survival is determined entirely by whether a cell’s lag time is sufficiently long to keep *λ*_avg_ below the critical threshold during the treatment period (*L* > *τ*).

This framework clarifies what makes transiently tolerant cells distinctive. Unlike persisters, which survive by remaining dormant beyond the treatment window (*L* > *τ*, rightward of the treatment boundary in Figure 7b), transiently tolerant cells decouple lag time from average growth rate: they wake early, with lag times comparable to the susceptible population, but their subsequent growth slowdown depresses *λ*_avg_, placing them in the lower-left region of Figure 7b (short lag, low *λ*_avg_, sublethal damage). This decoupling is a defining feature that separates transient tolerance from persistence as a survival mechanism. A small number of cells that follow the normal susceptible trajectory but happen to survive (for instance, because treatment duration is short or the concentration is low) occupy the boundary region near the critical *λ*_avg_; we refer to these as exposure-limited survivors (Box 2).

We formalised this conceptual separation as a state-transition model (Figure 7c; model structure and fitting detailed in Methods and Supplementary Note 3). The model tracks cells across three states during treatment: dormant (*D*), susceptible (*S*), and transiently tolerant (*T* ). Dormant cells resuscitate into *S* at rates governed by a heterogeneous lag-time distribution. Susceptible cells face concentration-dependent killing, and a fraction transition into state *T*, acquiring a reduced but non-zero death rate. The relationship between these model states and the experimentally observed post-treatment survivor classes is as follows (Figure 7c):

- Dormant cells that have not yet resuscitated by the end of treatment time (*τ*) (lag time *L* > *τ*) survive by remaining in state *D*. We refer to these as lag-based survivors. The extreme tail of the lag time distribution which contains these individuals corresponds to classical starvation-triggered persisters, which are a subset of the dormant pool (*P* ⊂ *D*).
- Cells that resuscitate during treatment (*L* < *τ*) enter the susceptible state *S*, where they face concentration-dependent rates of killing. A fraction of these susceptible cells transition into the transiently tolerant state *T*, acquiring a reduced but non-zero death rate; these are the transiently tolerant survivors whose *λ*_avg_ falls below the diagonal in Figure 7b.
- Susceptible cells that survive until and after drug removal without switching state (e.g. due to a short treatment time (*τ*), or low concentration (*C*)) are classified as exposure-limited survivors (Box 2) and thus a subset of class *S* (*X* ⊂ *S*); these are not considered a mechanistically distinct class, but rather a tail of susceptible cells whose growth trajectory in relation to treatment timing allowed them to escape killing. *Note:* Our model treats any cells that exit dormancy after the treatment window *τ* as moving from state *D* to state *P* (Figure 7c). As a result, otherwise susceptible cells which survive treatment due to very short treatment, get classified as *P*, not exposure limited (*X* ).

Thus, the three post-exposure survivor classes emerge naturally from the three model states. In the absence of the *T* state, the model reduces to the classical two-class system of rapid susceptible killing followed by a slow persister tail. Introducing *T* generates an intermediate effective death rate, producing a distinct shoulder in the survival curve before persister-dominated decay (Figure 7d).

To test whether this structure quantitatively reproduces the observed data, we fitted the model to the experimentally measured population fractions of each cell class across all tested combinations of antibiotic concentration, treatment duration, and culture age (Figure 8a–d). The model predicts the observed fractions across nearly three orders of magnitude in survivor frequency, spanning the full experimental range from sub-MIC to >15× MIC concentrations of ampicillin, from 2 to 8 hours of exposure, and from 24- to 72-hour-old cultures. The fraction of persisters (Figure 8d) is difficult to predict precisely, likely reflecting the very low cell counts in this class (typically <15 cells per condition, visible as large error bars in Figure 8d). Nonetheless, owing to the relative abundance of transiently tolerant versus persister cells, this deviation does not substantially affect the regrowth predictions (Supplementary Note 3).

**Figure 8:**
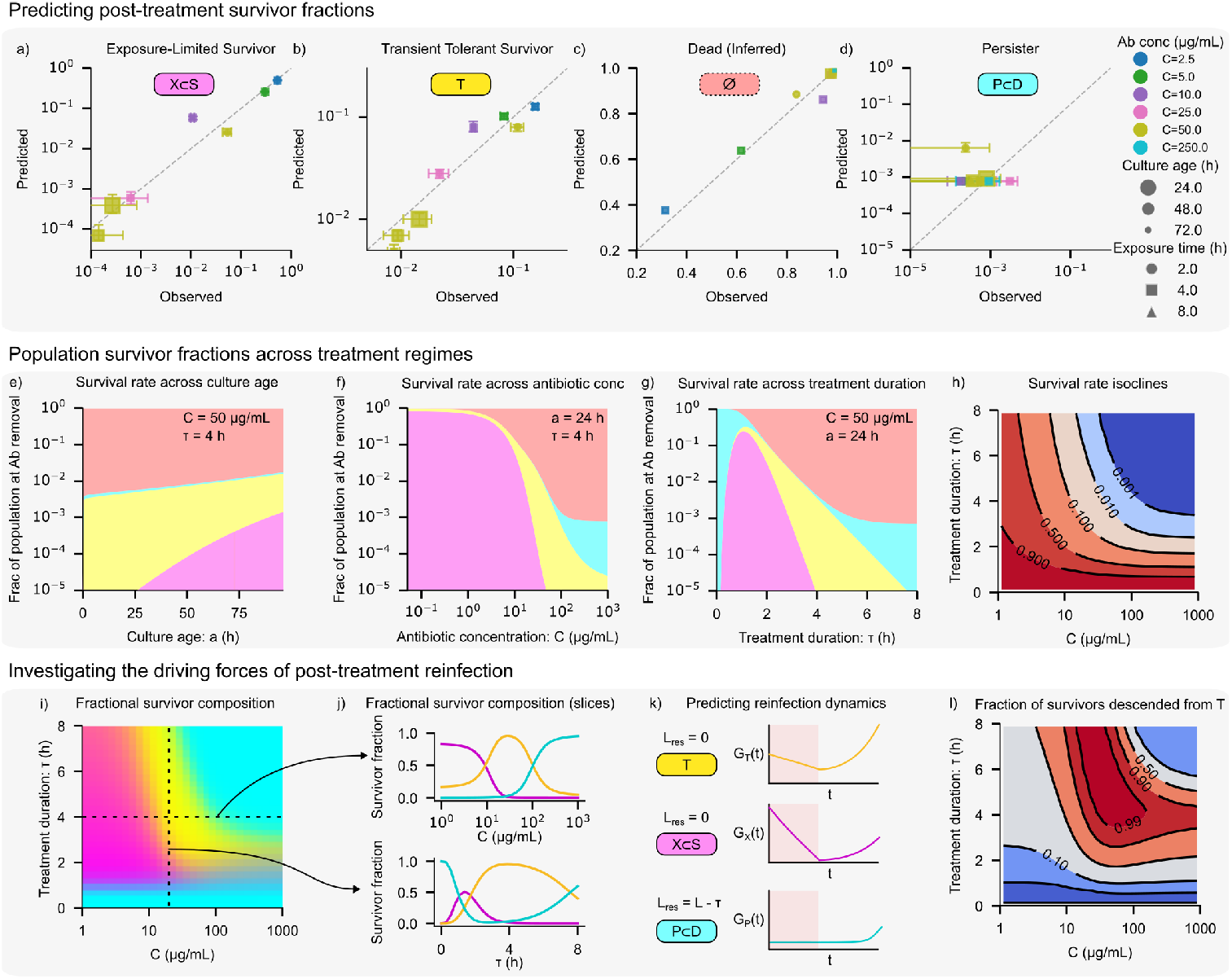
Transient tolerance dominates rapid population rebound under clinically relevant treatment conditions. **a–d)** Predicted versus observed population fractions for each cell fate class across all tested conditions: exposure-limited survivors (*X* ⊂ *S*, **a**), transiently tolerant survivors (*T*, **b**), dead cells (∅, inferred as 1 − *X* − *T* − *P* ; **c**), and persister/lag survivors (*P* ⊂ *D*, **d**). **e–g)** Model-predicted class distributions as a function of starvation history and treatment variables. **h)** Model-predicted total survival fraction as a function of treatment duration *τ* and antibiotic concentration *C*. **i)** Model-predicted fractional composition of the surviving population across the (*τ, C*) parameter space. **j)** Representative composition slices from panel i). **k)** Schematic post-treatment regrowth dynamics for each survivor class. **l)** Asymptotic fraction of the total regrowing population descended from transiently tolerant (*T* ) survivors as a function of concentration and treatment time.

With the model fitted against available experimental data from this study, we used it to map the full treatment parameter space and predict post-treatment outcomes (Figure 8e–h). The treatment outcomes are affected by culture age, with older cultures yielding progressively higher survival fractions, driven predominantly by transient tolerance and exposure-limited survivors (Figure 8e). The model also captures the characteristic crossover between survivor classes: exposure-limited survivors dominate at low concentrations or short treatment times, transiently tolerant cells are the majority survivor class at intermediate treatment conditions, and only at very high concentrations or prolonged treatment times do persisters become the dominant surviving class. The fractional composition of survivors shifts systematically across this landscape (Figure 8f and 8g): at short treatment times and moderate concentrations, exposure-limited survivors make up a substantial fraction, but their proportion declines steeply with increasing *τ* and *C*, giving way first to transiently tolerant survivors and then, at the most aggressive treatment regimes, to persister/lag survivors. The dominance of *T* across a broad intermediate region of the parameter space is a notable prediction of the model. The survival isoclines across treatment duration and concentration (Figure 8h) reveal a sharply non-linear landscape: survival drops by several orders of magnitude over a narrow band of the (*τ, C*) parameter space, implying that small changes in either treatment duration or concentration near this boundary can dramatically alter treatment outcomes.

Critically, however, the composition of the surviving population at drug removal does not directly predict which class drives relapse, because the three survivor classes differ fundamentally in their readiness to resume growth (Figure 8k). Both transiently tolerant and exposure-limited survivors have already exited dormancy during treatment and resume exponential growth immediately upon drug removal (residual lag *L*_res_ = 0). Persisters, by contrast, retain a residual lag of *L*_res_ = *L* − *τ*: they must first complete the remaining stages of their dormancy exit before contributing to the growing population. The model captures this asymmetry through a growth discount in the post-treatment regrowth simulations, whereby late-waking persisters produce exponentially fewer descendants than cells that begin dividing immediately (Supplementary Note 3). As a consequence, the fractional contribution of *T* -descended cells to the regrowing population (Figure 8l) substantially exceeds their fractional representation among survivors at drug removal (Figure 8i).

Across a wide region of the (*τ, C*) parameter space, spanning treatment durations of 2–6 hours and concentrations of ∼10–100 µg/mL ampicillin, as much as 90% of the early regrowing population descends from transiently tolerant cells (Figure 8l). This conclusion is robust to the underprediction of the persister fraction: across clinically relevant treatment conditions, transiently tolerant survivors outnumber persisters by 10–50-fold at drug removal, and the residual-lag penalty further amplifies this advantage during regrowth. The larger cell size of the *T* class at treatment end, associated with filamentation (Figure S7), may also amplify this effect further. Only when treatment durations are sufficiently long to substantially deplete the transiently tolerant subpopulation does the survivor composition shift toward deep persisters, whose delayed regrowth may demand a wider therapeutic window for subsequent dosing or immune-mediated clearance. This defines a practical criterion for therapeutic success with *β*-lactams: exposure must achieve the duration and concentration necessary to clear the transiently tolerant subpopulation, a requirement not captured by conventional metrics such as time above MIC.

## 3 Discussion

Antimicrobial resistance (AMR) remains a major global health threat, and heterotolerance[10], the ability of a subpopulation of bacteria to survive antibiotic exposure without acquiring genetic resistance, is increasingly recognised as a key contributor to treatment failure[45, 46]. Using our high-throughput dynamic fate analyser (Hi-DFA) pipeline, we identify a previously uncharacterised class of heterotolerant cells, which we term transiently tolerant cells. These transiently tolerant cells are highly prevalent under *β*-lactam treatment during resuscitation from starvation. Transiently tolerant cells exhibit a temporary period of slowed growth, modulated by prior starvation history, representing a dynamic form of heterotolerance that is fundamentally distinct from classical starvation-triggered persisters. Persisters survive through prolonged lag and deep growth arrest, while transiently tolerant cells resume growth but transiently modulate their growth rate in a manner that increases survival probability. While growth-rate fluctuations have been observed under *β*-lactam treatment of active cells in balanced growth[47], such fluctuations have been attributed to immediate antibiotic effects. In contrast, our work demonstrates that transient tolerance is programmed by prior stationary-phase stress and is highly prevalent during the resuscitation phase, revealing a distinct survival regime not captured in earlier analyses. Recent studies have also described metabolically mediated transient tolerance at the population level[48]. Our single-cell measurements suggest that, during resuscitation or nutrient switch, such population-level behavior may arise from enrichment of a discrete starvation-history-dependent heterotolerant subpopulation.

Culture age strongly modulates the prevalence of the transiently tolerant survivors. Extended duration in the stationary phase tends to increase the frequency of all survivor classes, but this effect is most pronounced for this transient growth slowdown class. Importantly, the stationary phase is not solely defined by nutrient depletion but is accompanied by multiple physiological stresses, including energetic limitation, redox imbalance, and accumulation of damage, which may collectively contribute to this stress-history dependent response. We used the term “starvation-primed” to describe this phenotype, since nutrient depletion is the initiating event that drives entry into stationary phase: secondary stresses such as accumulation of toxic metabolic by-products, oxidative damage, and pH changes are downstream consequences of growth arrest caused by carbon-source exhaustion[49–52]. Furthermore, the quantitative scaling of transiently tolerant survivor frequency with starvation duration (Figure 5) is most parsimoniously explained by a progressive, time-dependent process initiated by nutrient limitation rather than by an acute stress event. Mechanistically, this study does not assign molecular drivers underlying the starvation history dependent phenotype. However, our findings highlight the need to identify the regulatory and metabolic circuits that retain information about past stress and modulate antibiotic susceptibility upon recovery. The study posits that starvation is not a passive, static state, but one whose duration shapes the subsequent antibiotic response of the cell.

Our modeling and experimental data indicate that transient tolerance, not starvation-triggered persistence, is the immediate driver of population regrowth *in vitro*. The higher frequency and rapid recovery of transiently tolerant survivors make them a key threat to treatment success, underscoring the importance of understanding and targeting this transient state. The dependence of transiently tolerant cells and persister frequencies on starvation history has potential implications beyond laboratory settings. In clinical contexts, bacterial populations often experience fluctuating nutrient availability[13, 14] and host-associated stressors that could act analogously to starvation[53]. Indeed, physiological conditions such as inflammation, anxiety, or weight loss have been associated with altered nutrient absorption and increased nutritional stress in the gut microbiome[54]. Such stress-induced environments may prime bacterial populations towards these transiently tolerant states, potentially reducing antibiotic efficacy and contributing to infection relapse. The principles uncovered are likely not restricted to *β*-lactams. Other cell-wall-targeting antibiotics such as fosfomycin and vancomycin, which also rely on active growth for their bactericidal activity, may similarly reveal transient tolerance among stressed populations. Establishing the clinical relevance and prevalence of this phenotype will require systematic testing across diverse species and clinical isolates under infection-mimicking conditions.

Our findings also challenge classical PK/PD paradigms, which typically focus on metrics such as time above MIC[55]. Our integrated experimental-computational approach demonstrates that successful clearance of stress-primed, transiently-tolerant cells requires not only sufficient antibiotic exposure but also careful consideration of pharmacokinetic shape. Transiently tolerant survivors resuscitate rapidly after drug removal, and short or suboptimal treatments fail to eliminate them. Even prolonged low-concentration treatments or short high-concentration treatments can permit relapse, while sufficiently long treatments at concentrations several-fold above the MIC are required for near-complete eradication. By combining Hi-DFA with microfluidic simulations of clinically relevant pharmacokinetic profiles, we show that “time and concentration above a heterotolerance-clearing threshold” may be a more relevant metric than conventional “time above MIC”, highlighting the importance of population heterogeneity and the temporal nature of antibiotic treatment—both of which are masked by MIC values. Beyond conventional dosing strategies, our results also suggest the potential for complementary interventions, such as metabolic priming or adjuvants that transiently stimulate cellular activity, to disrupt starvation-history-dependent tolerance and thereby sensitize otherwise tolerant populations[56, 57]. Together, these directions underscore how understanding physiological history and transient tolerance can inform new approaches to antibiotic treatment design aimed at reducing relapse risk.

While this study focused on a single lab strain, a defined growth medium, and a limited set of antibiotics, these choices were deliberate to prioritise development and validation of the Hi-DFA platform. Hi-DFA enables high-throughput, time-resolved single-cell fate tracking, while the programmable microfluidic antibiotic treatment system allows simulation of clinically relevant pharmacokinetic profiles, and the computational model provides predictive insights that extend beyond the experimental dataset. Together, this study presents and establishes a quantitative foundation that extends beyond the specific experimental conditions presented here. A key consideration of single-cell microfluidics is the physical isolation of lineages, precluding direct cell-cell interactions. In bulk populations, such interactions (for example, the inoculum effect, where increased local cell density can shorten lag times and narrow their distribution[58]) may modulate regrowth dynamics after drug removal. However, this isolation is also a central strength of the Hi-DFA approach: in conventional bulk assays such as CFU counting, fast-growing survivors rapidly outcompete slow-resuscitating cells, obscuring the identity and relative contribution of each survivor class. By preventing competition between lineages, the MMX design allows us to directly resolve which phenotypic class survives and how quickly they resume growth, parameters that are inaccessible in the bulk. These lineage-resolved capabilities underpin the predictive capabilities of our population model. While cell-cell interactions may modulate the precise dynamics of population recovery *in vivo*, the lineage-resolved single-cell measurements underlying the model provide a level of phenotypic resolution that is necessary for quantitatively predicting survival dynamics, relapse risk, and pave the way to developing optimal treatment strategies in diverse bacterial contexts. These measurements isolate the intrinsic contributions of distinct survivor classes, which can subsequently be integrated with interaction-dependent effects to improve predictive models of relapse risk and treatment optimization across bacterial contexts.

Although this study was limited to *in vitro* conditions with *β*-lactam antibiotics on laboratory *E. coli* strains, the predictive framework established here can be extended to clinical isolates of multiple species, additional antibiotic classes, and more physiologically relevant media. The MMX trench architecture is directly compatible with the majority of high-priority pathogens: of the seven ESKAPE+*E* species, four (*Enterococcus faecium, Klebsiella pneumoniae, Enterobacter* spp., and *E. coli*) are rod- or coccal-shaped organisms whose dimensions permit reliable trapping in the current device[59–61]. While the dimensions of *Staphylococcus* cells seems permissible, the cluster-like growth pattern of this species prevents the use of the linear trench design. Species that grow and elongate along a single axis under *β*-lactam exposure, as *E. coli* or *Enterococci*, are particularly well suited to our growth-rate-based phenotyping pipeline. However, the current trench geometry does impose constraints: smaller organisms such as *Acinetobacter baumannii* may not be reliably retained, and species that swell isotropically rather than elongate (e.g., *A. baumannii* under *β*-lactam stress) would require modified trench widths and adapted morphological classifiers. These are engineering refinements rather than fundamental limitations of the Hi-DFA framework, and future iterations of the device will target expanded species compatibility.

In conclusion, by defining and mechanistically characterising a transiently tolerant subpopulation and integrating Hi-DFA, microfluidics, and predictive modeling into a single platform, this study delivers a novel conceptual and methodological approach to evaluate effective antibiotic treatment. It establishes a generalisable, predictive framework for studying bacterial response and recovery from treatment, reshapes how antibiotic efficacy can be assessed, and lays the groundwork for further informing treatment strategies, which could substantially reduce infection recurrence.

## 4. Methods

### 4.1 Chemicals and antibiotics

All chemicals and antibiotics were purchased from Sigma or Thermo Fisher Scientific. Stock solutions of ampicillin (50 mg/mL), cefalexin (10 mg/mL), and ceftriaxone (10 mg/mL) were prepared in distilled water. Stock solutions of amoxicillin (10 mg/mL) were prepared in distilled water and 10 M NaOH added dropwise until the powder was dissolved and pH was around 8.0. M9 minimal medium was prepared from 1× M9 salts, MgSO_4_ (2 mM), CaCl_2_ (0.1 mM), and trace elements solution (containing ZnSO_4_ (6.3 µM), CuCl_2_ (7 µM), MnSO_4_ (7.1 µM), CoCl_2_ (7.6 µM)) in distilled water.

### 4.2 Strains and culture conditions

*E. coli* MG1655 CGSC 6300[62] (“SB1”[63]) was used for all experiments. Cells were routinely cultured in supplemented M9 media, containing glucose and casamino acids, both to a final concentration of 0.05%, and pluronic F-108 to a final concentration of 0.08%. Overnight cultures were incubated at 37 °C, with shaking at 200 rpm.

Each individual experiment was performed using independent overnight cultures inoculated from frozen glycerol stocks stored at −80 °C. Overnight cultures inoculated from glycerol stocks were diluted 1:100 in fresh media exactly 24 hours before loading into the MMX device, with the exception of the different culture age experiments in Figure 5, where the cultures were diluted 1:100 either 24, 48, or 72 hours prior to cell loading. For the primary experiments (Figures 3–5), data were pooled across multiple independent device runs performed on separate days; the total cell numbers reported in each figure legend reflect the combined dataset. The high throughput of the MMX platform (up to 19,200 cells per condition per device) means that each run itself samples thousands of independent cell lineages, providing statistical power comparable to many biological replicates in conventional assays. Where error bars are shown, they represent 95% confidence intervals calculated by bootstrapping (*n* = 10,000).

### 4.3 Antimicrobial susceptibility testing

Minimum inhibitory concentrations (MICs) were determined by broth microdilution. Briefly, overnight cultures in supplemented M9 media were diluted 1,000× into fresh media. 50 µL were added to wells of flat-bottom 96-well microtiter plates already containing 50 µL of 2-fold serially diluted antimicrobials in supplemented M9 media, to a final volume of 100 µL. The plates were sealed with a gas-permeable seal and incubated at 37 °C for 18 hours. Growth was determined by visual inspection and OD_600_ measurement using a SpectraMax iD3 microplate reader.

### 4.4 Microfluidic device fabrication and cell loading

Microfluidic devices were fabricated as described in[64]. Briefly, PDMS for casting the device (Sylgard 184, Dow) was prepared by mixing base elastomer and curing agent at a 10:1 (w/w) ratio, degassed under vacuum, poured over an SU-8 master with appropriate features for the MMX device, and cured at 95 °C for 1 h. The cured PDMS was peeled off, devices were cut individually, and inlet/outlet ports were created using a 0.75 mm biopsy punch. PDMS replicas were cleaned by sequential washing with isopropanol and deionised (DI) water, with intermediate drying and baking at 95 °C. Glass coverslips were cleaned by washing in 1 M KOH followed by DI water, then dried and baked. PDMS devices were plasma-bonded (35 W, 2 min, 0.2–0.3 mbar) to glass coverslips (feature side down) and baked at 95 °C for 30 min to strengthen the bond. Bond integrity was verified by stereo microscopy prior to cell loading. For cell loading, stationary phase cultures of cells were centrifuged at 1,000 × *g* for 3 minutes, resuspended in approximately 5% of the initial volume, then 5–10 µL was added to each lane using long gel loading tips.

### 4.5 Timelapse microscopy and treatment

Timelapse microscopy of cells loaded in the MMX microfluidic device was set up as described in[65], using a Nikon Eclipse Ti2 inverted microscope with a Hamamatsu ORCA-Fusion Digital CMOS camera, with a pixel size of 6.5 µm × 6.5 µm. Multiple lanes of the MMX device were imaged in parallel using an automated *xy* stage and Nikon Perfect Focus System to maintain focus throughout the experiment. The device was maintained at 37 °C in a custom enclosure. All acquisitions were carried out using a 40× 0.95 NA objective lens, with images taken in phase contrast every 3 minutes, with the exception of the ampicillin supra-MIC survival experiment used in Figure 4c,e, which was carried out using a 20× 0.75 NA objective lens (with 1.5× post-magnification) with images taken every 70 seconds. Phase contrast images were taken with 30 millisecond exposure per frame and 80% LED power. For pharmacokinetics treatment experiments, the cells were also imaged in the GFP channel (ex: 488 nm, em: 515/30 nm), illuminated by a Lumencor Spectra III fluorescence light source every 3 minutes with 200 millisecond exposure per frame and 50% LED power.

After cell loading and the start of image acquisition, filter-sterilised spent media from the overnight cultures flowed through the device, typically for 30 minutes, followed by 2–8 hours (typically 4 hours) of fresh supplemented M9 media containing the specified concentration of antibiotic. The same supplemented M9 media without any antibiotic was then flowed for typically 16 hours. A NE-300 syringe pump was used. Flowrates of 5 µL/minute were used throughout, with temporary increases to 50 µL/minute for 5 minutes during media switches.

### 4.6 Pharmacokinetics treatment experiments

For experiments using the pharmacokinetics (PK) experiments, microfluidic devices were first loaded with cells, and then spent media was flowed through the device. The PK setup was also primed with spent media. The primed PK setup was then connected to the inlets of the microfluidic device (detailed in Supplementary Note 2). A flow sensor was attached to the outlet of the microfluidic device and a pump speed calibration was run to determine the required pump speed for the desired flowrate. Following calibration, the pumps were stopped and the spent media in the PK setup was removed and replaced with fresh media. Tubes were connected containing either fresh media or fresh media + concentrated antibiotic + fluorescein. After restarting the pumps, the concentration of antibiotic in the microfluidic device was measured by green fluorescence intensity (of the fluorescein) of the feeding lane of the device. The fluorescein dynamics of the feeding lane were consistent with the dynamics within individual trenches (Figure S9). At the end of each experiment, a known dilution of the same fresh media + concentrated antibiotic + fluorescein stock was flowed through the device to calibrate the fluorescence intensity values obtained during the experiment.

Prior to an experiment with the PK setup, the tubes and tubing were sterilised with approximately 5 mL of 10% bleach, followed by 5 mL of 70% ethanol, followed by 10 mL of sterile water.

### 4.7 Image processing and single-cell data analysis using Hi-DFA

Image data were saved, registered, and extracted to images of single trenches using custom-designed image-processing pipelines[29, 66]. Individual cells were identified from single-trench images using Omnipose[25, 67] with models initially trained on synthetic images generated by SyMBac[26, 68] and retrained on manually curated real images. Custom Python scripts were used to extract properties including area, length, and width for the top cell of each trench for further analysis as described in Supplementary Note 1. The Hi-DFA pipeline is available at https://github.com/kieranrabbott/Hi-DFA.

### 4.8 Statistical analysis

Unless otherwise stated, data are presented as means with error bars indicating the upper and lower 95% confidence intervals from bootstrapping with *n* = 10,000. The number of cells comprising the data are given in the corresponding figure captions. Pre-processing of the cell length data was carried out within the Hi-DFA pipeline as described in Supplementary Note 1.

### 4.9 Modelling treatment and post-treatment relapse dynamics

To predict which survivor class drives post-treatment population rebound at different treatment regimes, we constructed a phenomenological population dynamics model that tracks single-cell states and associated fates during and after antibiotic exposure. The model describes transitions among three cellular states: dormant cells that have not yet resuscitated (*D*), susceptible actively growing cells (*S*), and transiently tolerant cells (*T* ). The correspondence between these model states and the experimentally observed survivor phenotypes is summarised below in Table 1.

**Table 1:** Mapping between experimental phenotypes and model states.

| Experimental term | Definition | Model state(s) |
| --- | --- | --- |
| Lag-only survivor | Cell does not exit dormancy within $\tau$ ($L > \tau$) | $D$ |
| Starvation-triggered persister | Lag-only survivor with exceptionally long growth arrest | $P \subset D$ |
| Susceptible cell | Cells that wake under the drug, and do not switch state | $S$ |
| Exposure-limited survivor | Susceptible cells that happen to survive treatment | $X \subset S$ |
| Transiently tolerant survivor | Wakes, transiently slows growth | $T$ |

Crucially, in this model, **starvation-triggered persisters are not a separate compartment**. Instead, they correspond to the far right tail of the lag distribution within the dormant state *D*; they are a *subset* of lag-only survivors.

The population dynamics during antibiotic treatment at constant concentration *C* over the window [0, *τ*] are governed by

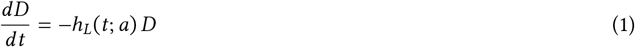

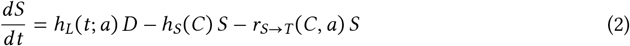

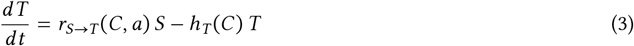

where *h*_*L*_(*t*; *a*) is the instantaneous hazard of exiting dormancy at time *t*, given culture age *a*. We assume lag time is independent of the antibiotic concentration. During treatment, susceptible cells can either die at rate *h*_*S*_(*C*) or switch to the transiently tolerant state at rate *r*_*S*→*T*_ (*C, a*). Cells that have become transiently tolerant die at rate *h*_*T*_ (*C*) < *h*_*S*_(*C*), so that tolerance reduces but does not eliminate the risk of death. Cells that die from either *S* or *T* enter an absorbing dead state.

Each rate is specified as follows:

#### Dormancy exit rate

Each dormant cell draws a lag time *L* before resuscitation. We model the lag-time distribution as a two-component mixture:

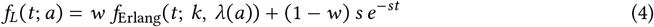

The primary component is an Erlang distribution (weight *w*, shape *k*, age-dependent rate *λ*(*a*)), which provides a biologically interpretable model of lag as a completion of *k* sequential events that admits a *k*-stage linear chain representation that preserves a Markov structure[69, 70] (Supplementary Note 3). The secondary component is a slow exponential (weight 1 − *w*, rate *s*) which captures a subpopulation of cells with extended lag times that are not well described by the Erlang tail. The mean of the Erlang component increases with culture age *a* (hours in stationary phase):

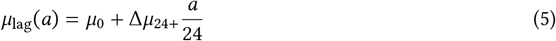

with Erlang rate *λ*(*a*) = *k*/*μ*_lag_(*a*), where *μ*_0_ is the baseline mean lag for a fresh culture and Δ*μ*_24+_ is the additional mean lag accumulated per 24 hours of stationary phase. The Erlang shape *k* (selected by grid search over *k* = 2, …, 14) determines the regularity of wake times: larger *k* produces a sharper distribution around the mean (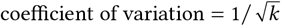). The overall mean lag time under the mixture is *E*[*L*] = *w μ*_lag_(*a*) + (1 − *w*)/*s*.

Rather than tracking the full lag-time distribution explicitly, we describe exit from dormancy by a **lag hazard** *h*_*L*_(*t*; *a*), the instantaneous rate at which dormant cells wake at time *t*, given that they have not yet woken.

#### State switching rate

The rate at which susceptible cells switch to the transient tolerant state is

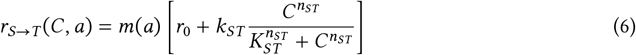

where *m*(*a*) = *a*/(*a* + *a*_50_) is a starvation-history modulation function that increases monotonically with culture age *a*, capturing the observation that older cultures produce more tolerant survivors (Figure 5b). The baseline rate *r*_0_ > 0 accounts for transient growth slowdown events observed even in untreated resuscitating cells (Figure S5), while the Hill term captures the concentration-dependent potentiation of switching. Note that we do not fit *r*_0_, but rather calculate it directly from our observation that ∼5% of cells exhibit a transient tolerant-type slowdown behaviour in the absence of antibiotics.

#### Concentration-dependent cell death hazards

Death rates for both susceptible and tolerant cells follow Hill kinetics sharing a common Hill coefficient *n* and half-maximal concentration *K* :

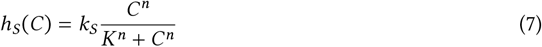

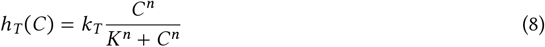

where *k*_*S*_ > *k*_*T*_ is a constraint imposed on the model, and we define *k*_*S*_ ≡ *k*_*S*/*T*_ *k*_*T*_ . Thus *h*_*S*_ > *h*_*T*_ at all concentrations. Sharing *n* and *K* between the two hazards encodes our assumption that tolerance acts by reducing the maximal kill rate rather than by shifting the dose-response curve[10].

#### Physiological constraints

We impose the constraint *λ*_eff_(*a*) ≤ *k*_*T*_ for all culture ages in the dataset, where *λ*_eff_(*a*) = 1/*E*[*L*] is the effective lag rate accounting for both mixture components. This ensures that at saturating antibiotic concentrations, the transiently tolerant kill rate (*h*_*T*_ → *k*_*T*_ ) equals or exceeds the effective rate at which dormant cells enter the susceptible pool, so that extended high-dose treatment selects for deep persisters rather than transiently tolerant survivors, consistent with our observation of complete killing at 250 µg/mL (Figure 4e). Using *λ*_eff_ rather than the Erlang rate *λ* is necessary because when the slow-exponential component dominates (small *w*), the actual wake rate is governed by *s*, not *λ*. Details of the constraint implementation are provided in Supplementary Note 3.

#### Survivor fractions

Because *h*_*S*_, *h*_*T*_, and *r*_*S*→*T*_ are constant during the treatment window (constant *C* and *a*), the *S*/*T* dynamics reduce to a linear system with constant coefficients driven by the timevarying inflow from *D*. This permits closed-form analytical expressions for the population fractions of each survivor class at time *τ*, obtained by convolution of single-cell survival kernels with the hybrid lag-time density (Supplementary Note 3). The model has 13 free parameters (Table 2), fitted by maximum likelihood using a Dirichlet scoring objective with multistart optimisation (*n* = 240 random initialisations). Sensitivity of the fitted parameters and model predictions is assessed via Laplace’s-approximation marginal distributions and sensitivity analysis of model outputs to parameter fluctuations (Supplementary Note 3).

**Table 2:** Model parameters for the Hill-Hybrid model (13 free parameters). The Erlang shape *k* is selected by grid search over *k* = 2, …, 14; all other parameters are fitted by L-BFGS-B multistart optimisation. Note that *g* is fixed, used only for regrowth simulations, and is not utilised in model fitting, and thus is not counted as a parameter.

| Symbol | Biological meaning | Fitted | Value |
| --- | --- | --- | --- |
| $k$ | Erlang shape parameter (number of lag stages) | Yes | 6 |
| $w$ | Erlang mixture weight ($w = 1$: pure Erlang) | Yes | 0.999 |
| $s$ | Slow-exit rate, exponential component ($\text{h}^{-1}$) | Yes | $1.63 \times 10^{-4}$ |
| $\mu_0$ | Baseline mean lag time (h) | Yes | 0.715 |
| $\Delta\mu_{24+}$ | Additional mean lag per 24 h of starvation (h) | Yes | 0.184 |
| $k_T$ | Transiently tolerant-state maximal kill rate ($\text{h}^{-1}$) | Yes | 3.55 |
| $k_{S/T}$ | Susceptible-to-tolerant kill-rate ratio ($k_{S/T} > 1$) | Yes | 2.26 |
| $k_{ST}$ | Maximal $S \rightarrow T$ switching rate ($\text{h}^{-1}$) | Yes | 1.74 |
| $K$ | Shared half-maximal concentration for $h_S, h_T$ ($\mu\text{g mL}^{-1}$) | Yes | 65.7 |
| $K_{ST}$ | Half-maximal concentration for $r_{S \rightarrow T}$ ($\mu\text{g mL}^{-1}$) | Yes | 32.6 |
| $n$ | Shared Hill coefficient for $h_S, h_T$ | Yes | 1.19 |
| $n_{ST}$ | Hill coefficient for $r_{S \rightarrow T}$ | Yes | 2.36 |
| $a_{50}$ | Culture age at half-maximal stress history (h) | Yes | 26.0 |
| $r_0$ | Baseline $S \rightarrow T$ switching rate ($\text{h}^{-1}$) | Calculated from data | 0.0608 |
| $g$ | Post-treatment balanced growth rate ($\text{h}^{-1}$) | Fixed | 1.2 |

#### Post-treatment regrowth

Upon removal of the antibiotic at *t* = *τ*, we model the competition among the three survivor classes to determine which drives the earliest population rebound. Let *u* = *t* − *τ* denote the time since drug removal. Exposure-limited (*S*) and tolerant (*T* ) survivors have already exited dormancy and resume exponential growth immediately:

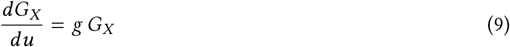

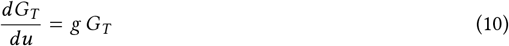

with *G*_*X*_ (0) = *N*_0_ *S*(*τ*) and *G*_*T*_ (0) = *N*_0_ *T* (*τ*), and where *g* is the balanced growth rate.

Lag-only survivors, such as persister cells, however, must first complete their residual dormancy period before exiting and contributing to the growing population. Their descendants are recruited from the dormant pool according to the residual lag distribution:

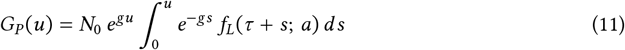

where *f*_*L*_(*τ* + *s*; *a*) is the hybrid lag-time density evaluated at *τ* + *s*, representing the rate at which dormant cells complete their residual lag at post-treatment time *s*. Both the Erlang and slow-exponential components contribute independently to this integral, with closed-form solutions available for each (Supplementary Note 3). We assume that these cells have remained unaffected during the entire period of treatment due to their lag.

The fractional contribution of each survivor class to the total population *N* (*u*) = *G*_*X*_ (*u*) + *G*_*T*_ (*u*) + *G*_*P*_ (*u*) then determines which class dominates population rebound at any post-treatment time.

The full model derivation, including the analytical survivor-fraction expressions, the post-treatment regrowth dynamics, the fitting procedure, and all parameter constraints, is given in Supplementary Note 3, where sensitivity analyses and comparisons with a pure-Erlang variant (11 free parameters) and a minimal linear-hazard model (7 free parameters) are also presented to explore model complexity and overfitting.

## Author Contributions

S.B. conceived the study and was in charge of the overall direction and planning. S.B. and A.Z. supervised the study. K.A., G.H., R.L., J.B., S.B., designed the experiments. K.A. performed microfluidic imaging experiments. K.A. and G.H. performed the data analysis. R.L. and S.B. designed the MMX microfluidic device. R.L. fabricated the MMX microfluidic device. J.B. and S.B. designed the pharmacokinetics hardware. J.B. built the pharmacokinetics hardware. K.A. and J.B. performed the pharmacokinetics experiments. G.H. developed the model, and performed the simulations and corresponding analysis. K.A., G.H., S.B., led the manuscript writing. All authors contributed to reviewing the manuscript.

## Acknowledgements

The authors thank Dr Antoine Hocher for their helpful comments and discussions.

## Supporting Information

### Supplementary Figures

**Figure S1:**
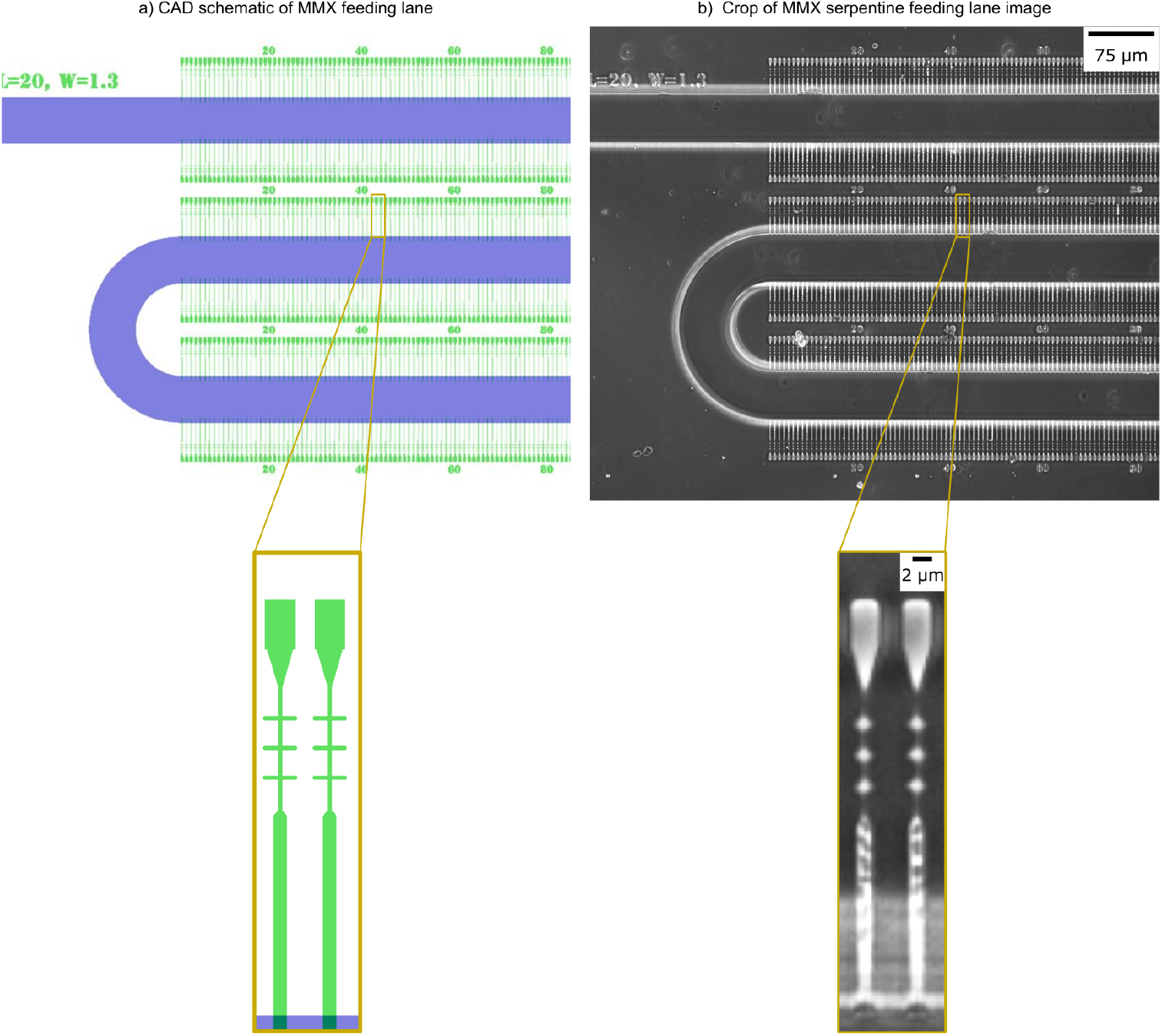
MMX flow lane and trench layout for high-throughput uniform treatment per lane. **a)** A CAD schematic showing the feeding lane (blue) and corresponding cell trenches (green), along with fiducial markers which allow us to identify the relative position of each trench in an FOV without knowing the absolute position of the microscope stage. The thickness of the feeding lane is 25 µm and 1.1 µm for cell trenches. **b)** A phase contrast image showing the PDMS cast MMX feeding lane and a zoom is shown below where bacterial cells are loaded in the trenches. The trench width is 1.3 µm, length is 20 µm, backport length is 12.5 µm, and width 0.4 µm, except the junctions. Finally, the evaporation chamber area is approximately 20.7 µm^2^. The typical snake-like layout of flow lanes lead to weak loading of regular mother machine trench design, possibly related to evaporation from parallel lanes through the separating PDMS affecting each other. The new trench design with evaporation chambers in MMX significantly improves loading in such scenarios.

**Figure S2:**
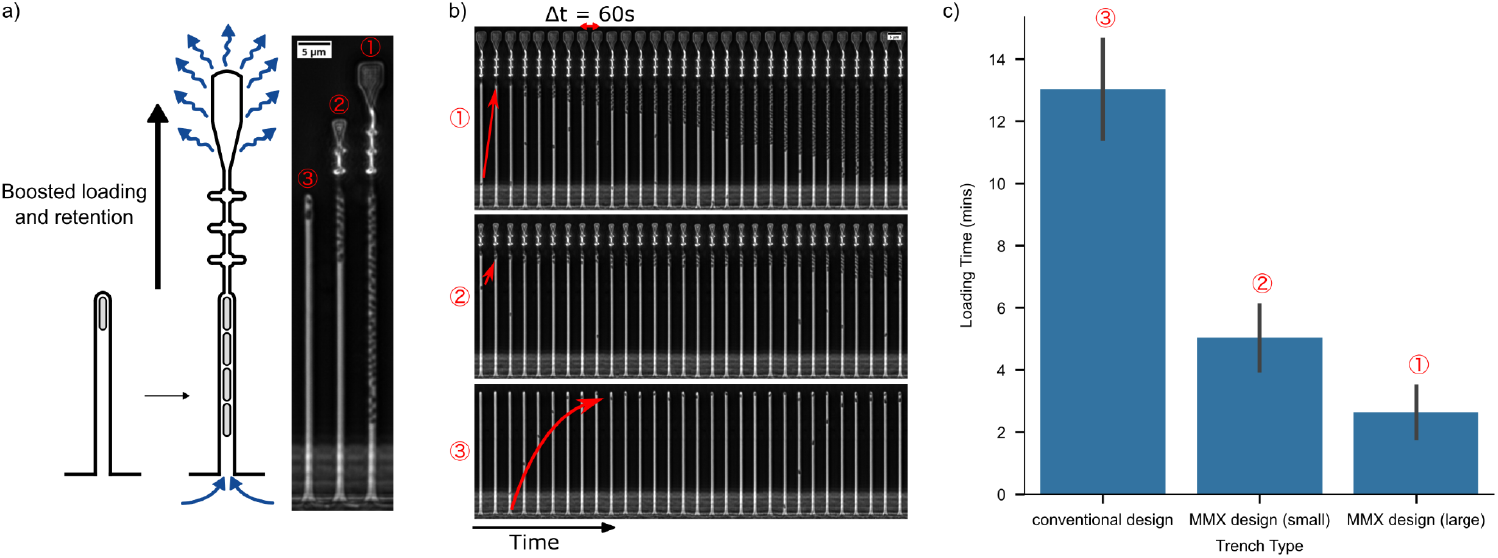
MMX trench design for evaporation-assisted efficient loading. **a)** The MMX trench design for evaporation assists efficient loading. A schematic cartoon is shown on the left. The real image (right) shows different designs benchmarked on the same test device and the same flow lane for direct comparison. **b)** Time-lapse imaging over a 30-minute period comparing loading dynamics in a MMX-design trench design (top), a MMX-design trench design with a smaller top chamber (middle), and a conventional mother machine trench (bottom). **c)** The bar plot shows the quantification of loading performance. Mean waiting times to obtain at least one cell per trench are 2.6 min (*n* = 40) for the MMX design, achieving ∼100% occupancy within 30 min.

**Figure S3:**
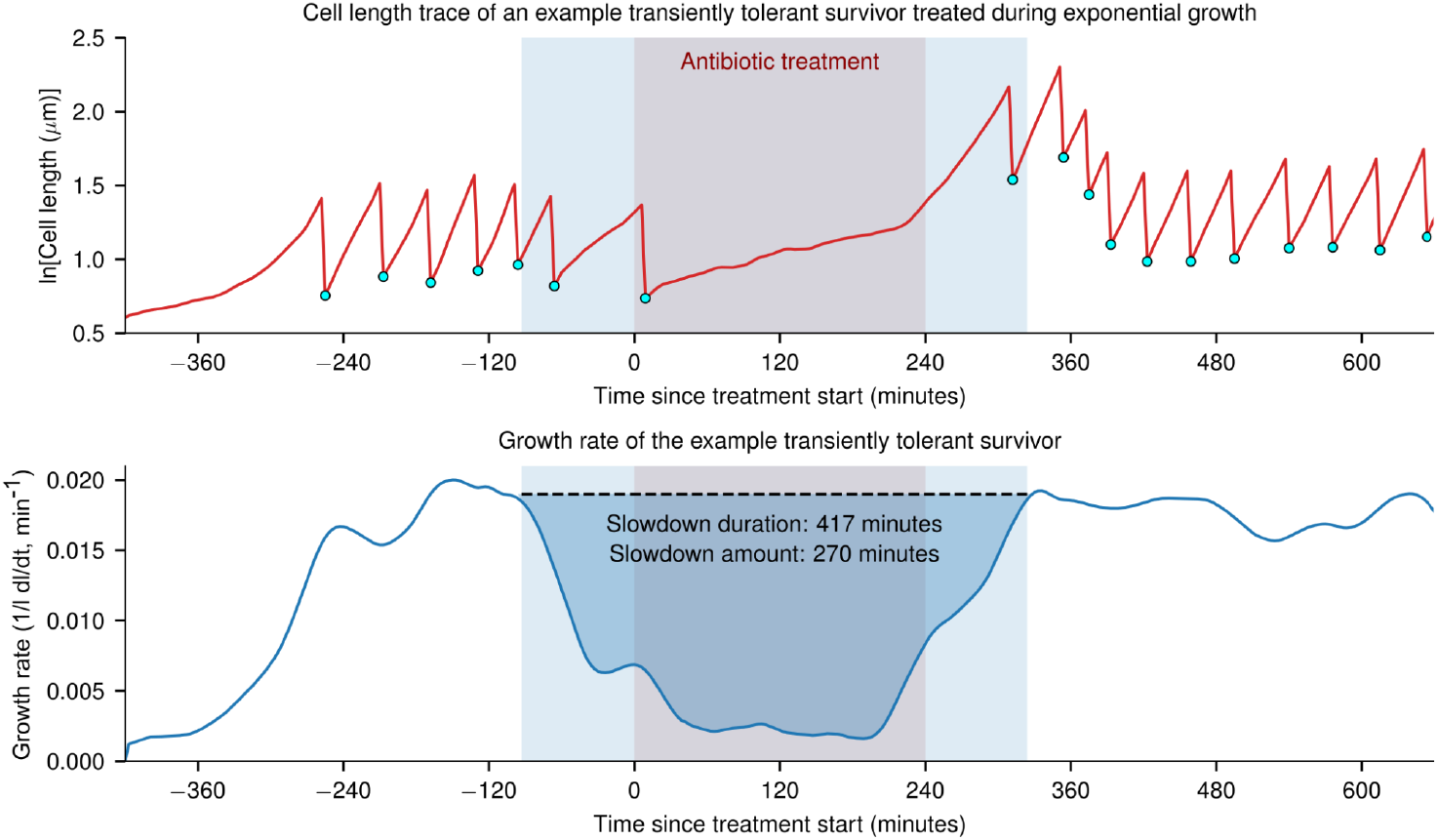
Exponential phase cells survive antibiotic exposure by spontaneous transient growth slowdown, not complete growth arrest. Plot of cell length against time (top) and corresponding plot of growth rate against time (bottom) for an example *E. coli* cell within an exponential phase population that survived 4 hours of treatment with 50 µg/mL ampicillin. A 24 hour old culture of cells was loaded into lanes of the MMX device and grown with fresh media for 7 hours prior to treatment start. Cyan circles indicate division events. Antibiotic treatment time is highlighted in red. The cell growth rate slowdown event is highlighted in blue. Slowdown duration corresponds to the time between the start and the end of the growth rate slowdown event. Slowdown amount refers to the total lag time accumulated during the slowdown event, using a similar definition to lag time in[71].

**Figure S4:**
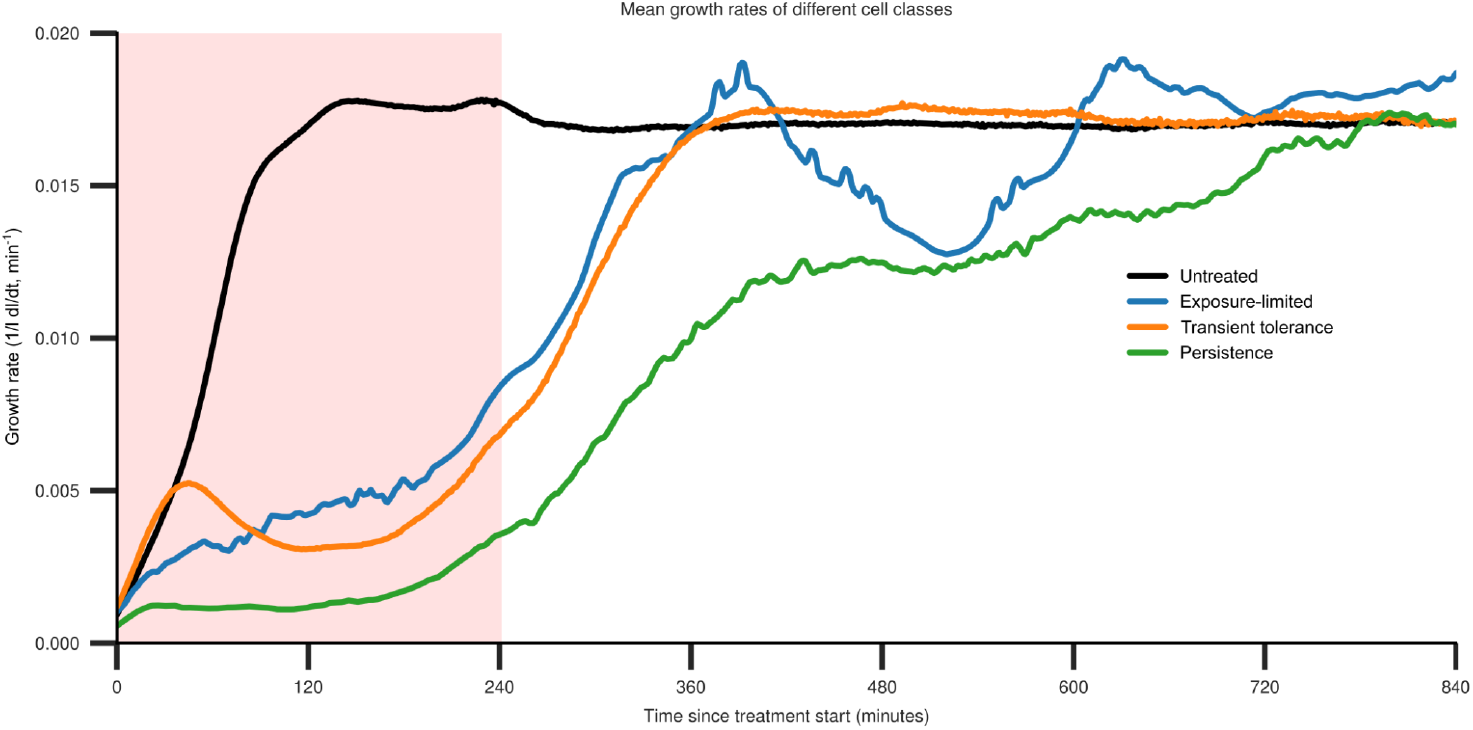
Mean growth rate of different cell classes. Black line indicates untreated cells resuscitated in the absence of an antibiotic. The transient tolerance class starts to follow the same trend as untreated cells or susceptible class, but then slows down growth during treatment and eventually reaches the maximal growth rate after the removal of treatment. Persister cells maintain slow growth throughout treatment and slowly catch up (depending on the distribution of lag time). Exposure-limited survivors are also caused by distributed lag time of the susceptible population and their mean growth rate increases slowly, relative to the susceptible population. Resuscitation was in the presence of no antibiotic (untreated) or 25 or 50 µg/mL ampicillin for 4 hours (red shaded region).

**Figure S5:**
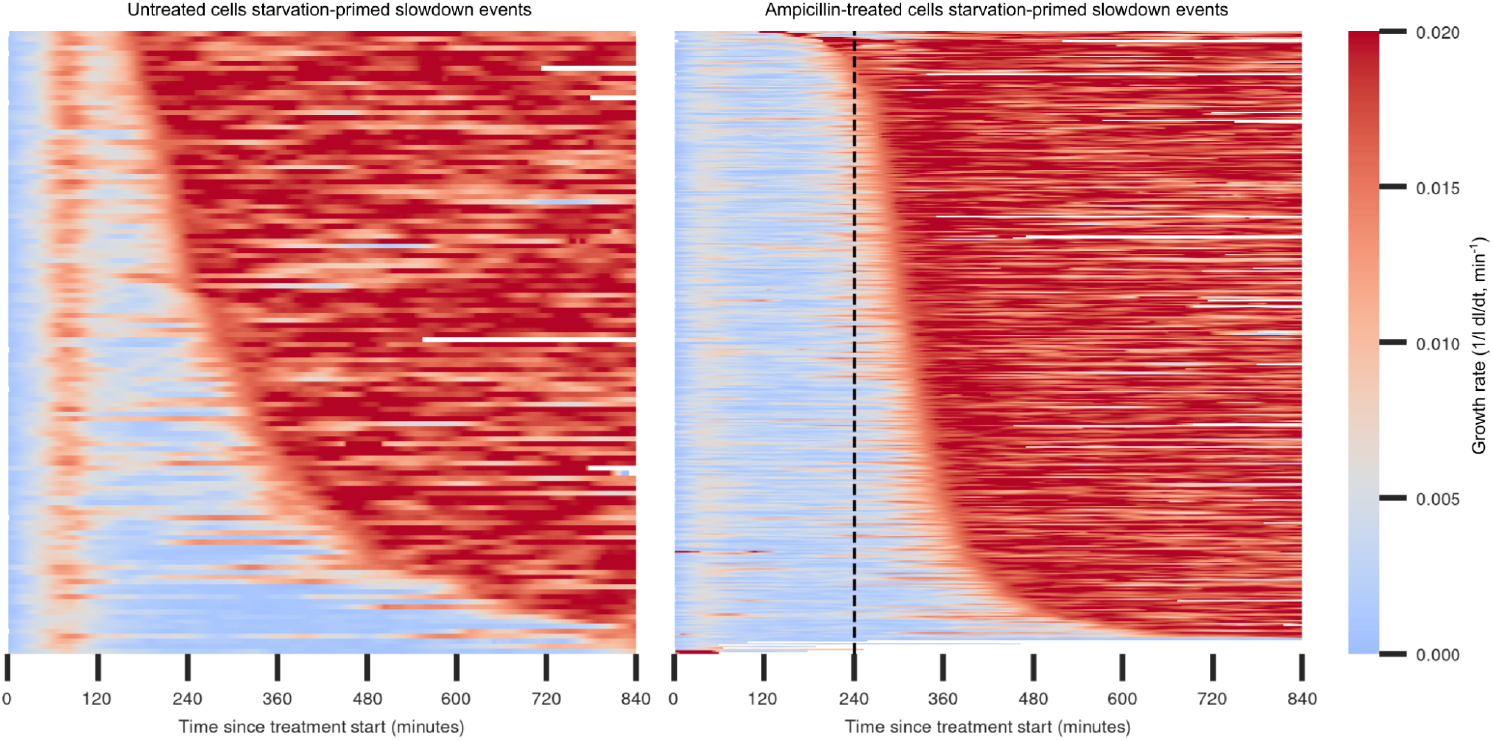
Transient growth slowdown events are present and heterogeneous in the untreated population (left). Heat maps of growth rates of slowdown events in untreated cells compared to the transiently tolerant subpopulation during and after treatment (right). The vertical dashed black line indicates the end of 4 hours of treatment with 25 or 50 µg/mL ampicillin for the transiently tolerant subpopulation.

**Figure S6:**
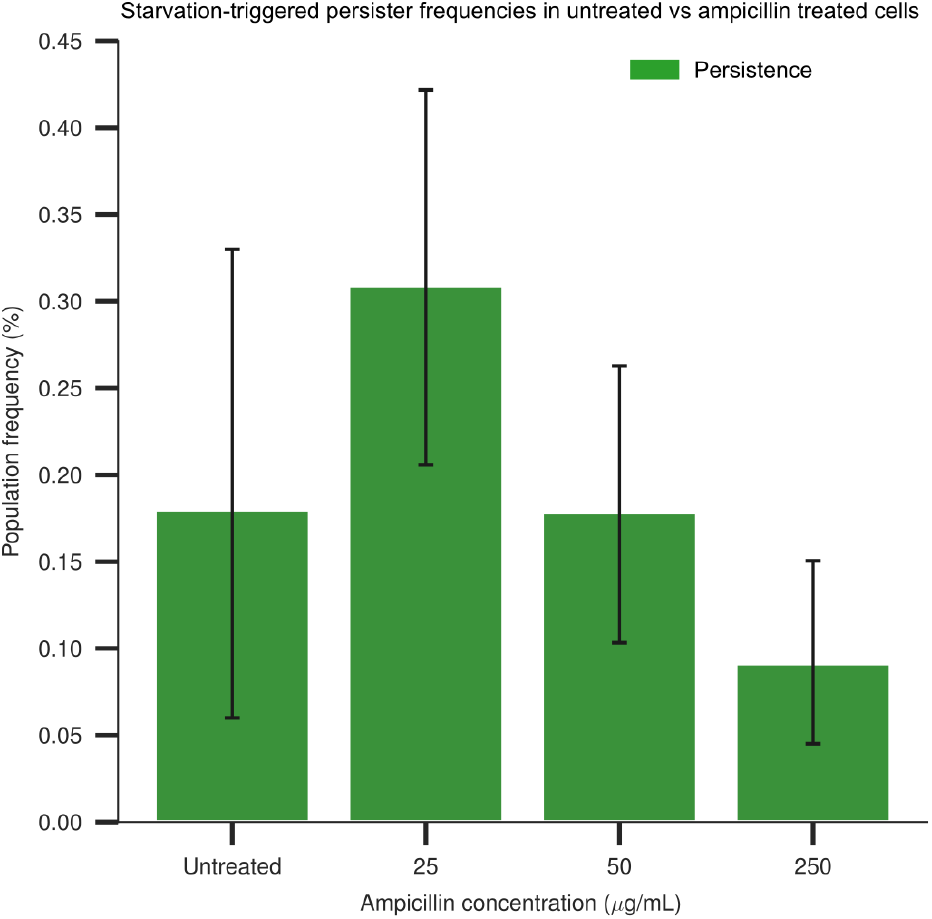
Starvation-triggered persister survivor frequencies of an untreated population of 24 hours old, compared to populations treated with varying concentrations of ampicillin for 4 hours. The initial frequency of starvation-triggered persisters in the population appears to be independent of antibiotic treatment, and hence instead depends on factors such as culture age. Error bars indicate the upper and lower 95% confidence intervals from bootstrapping with *n* = 10,000.

**Figure S7:**
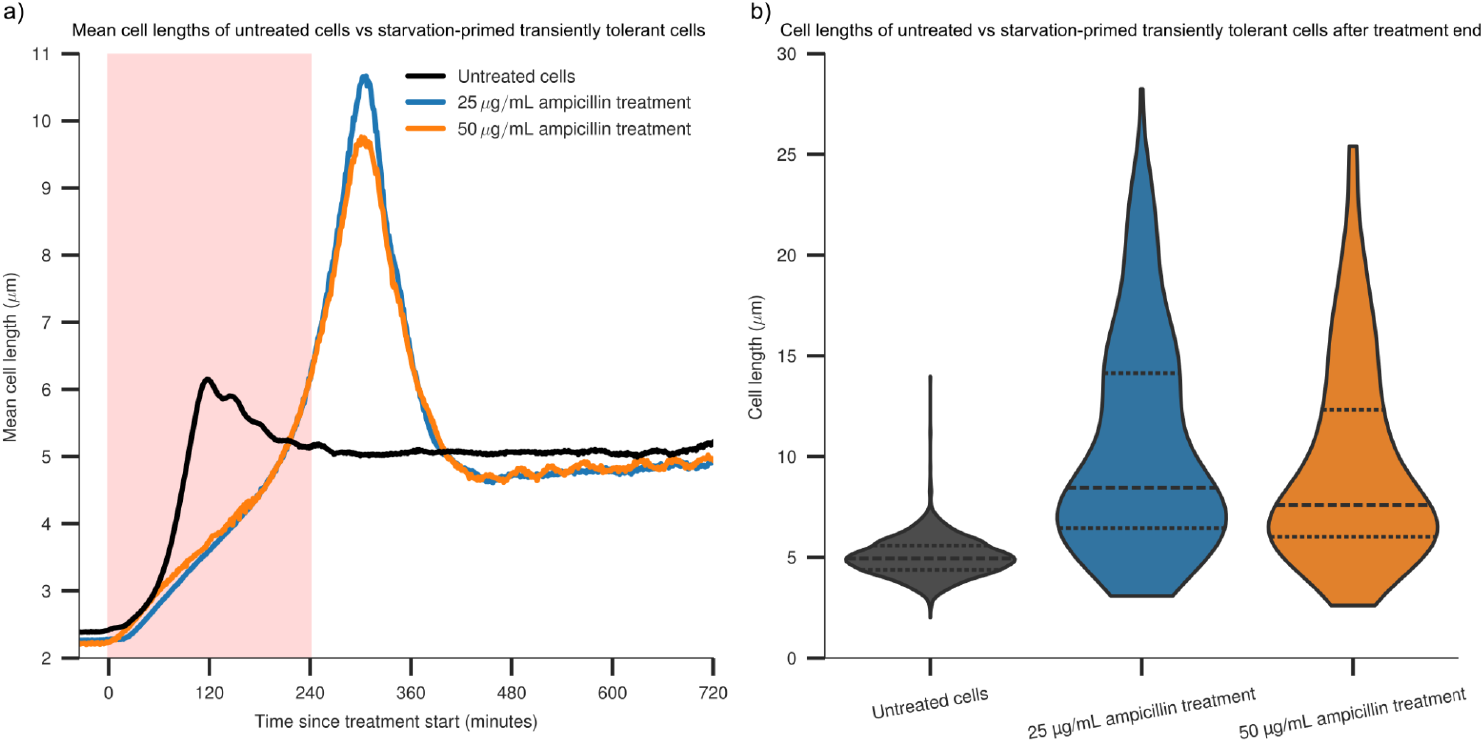
Transiently tolerant cells frequently filament to form long cells that can divide to a large number of progeny following treatment end. **a)** The mean value of single-cell length traces of a culture of 24 hour old *E. coli* resuscitated in the presence of different treatment conditions. The red shaded region indicates antibiotic treatment. **b)** Violin plots showing the distribution of single-cell lengths 60 minutes after the end of antibiotic treatment (300 minutes after treatment start).

**Figure S8:**
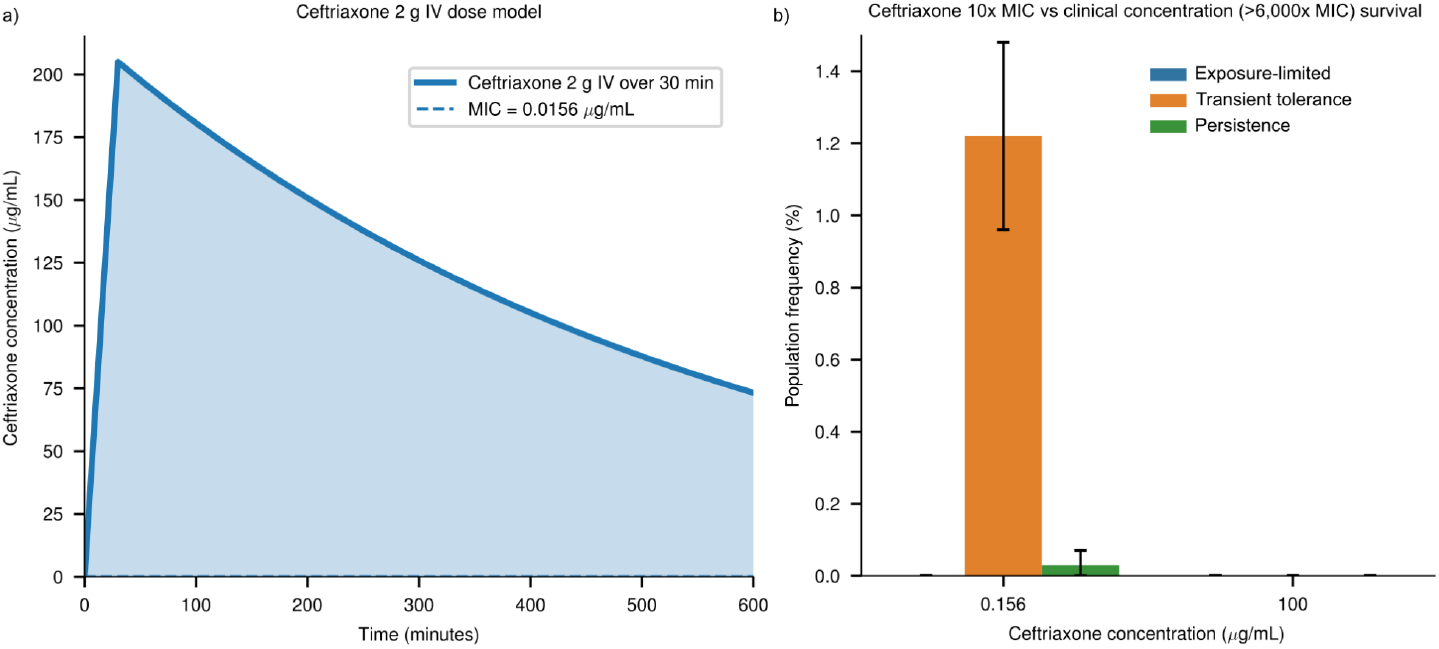
Ceftriaxone clinical pharmacokinetics model and survival at different concentrations. **a)** Pharmacokinetics model of a typical 2 g dose of ceftriaxone administered by IV over 30 minutes. Based on PK parameters from[40, 41]. **b)** Survival frequencies of 24 hour old cultures of *E. coli* treated with either 10× MIC (0.156 µg/mL) or the more clinically relevant >6,000× MIC (100 µg/mL) of ceftriaxone for 4 hours. Error bars indicate the upper and lower 95% confidence intervals from bootstrapping with *n* = 10,000.

**Figure S9:**
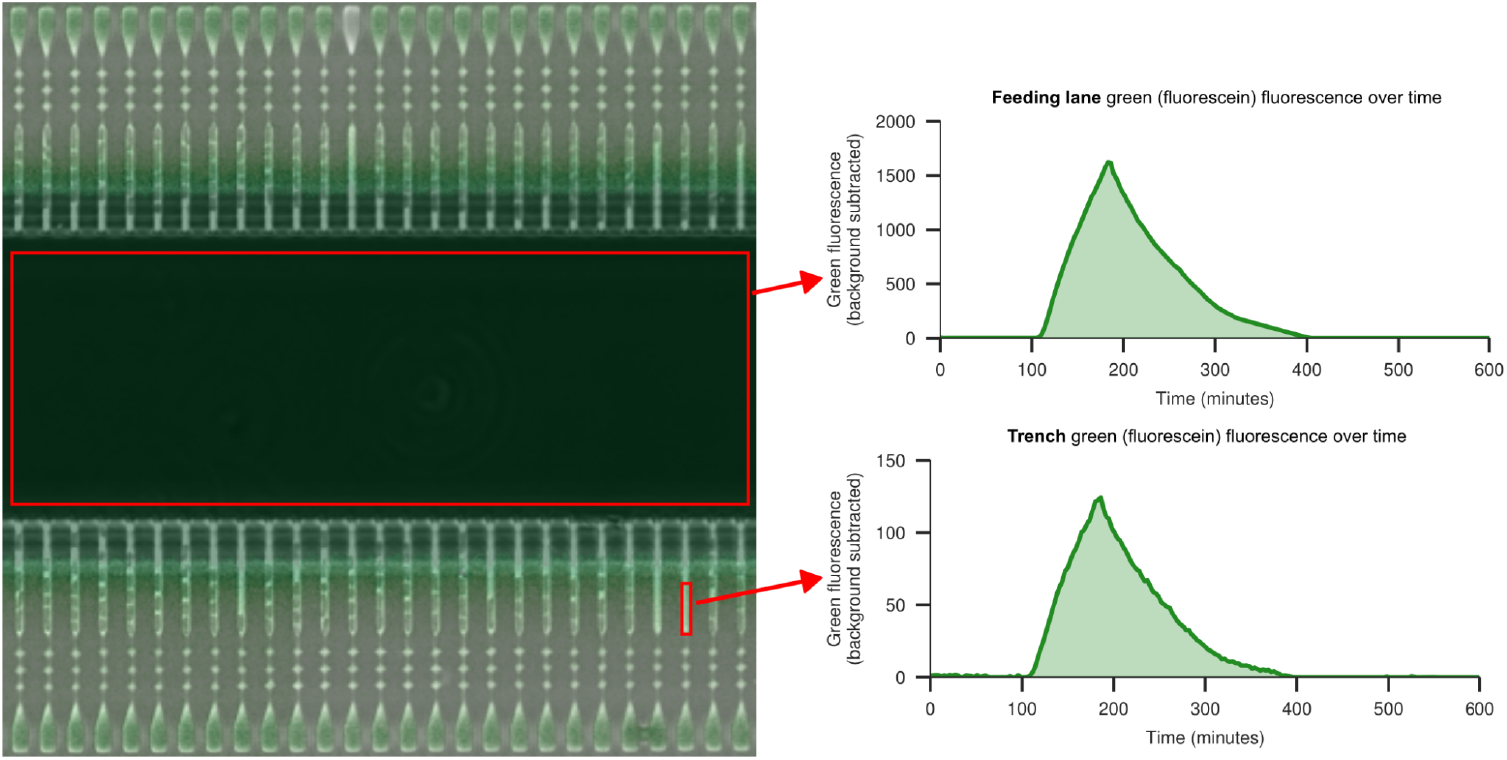
Fluorescein dynamics are consistent between the feeding lane and individual trenches. Overlayed image of the green fluorescence channel and the phase contrast channel of a section of the MMX device 180 minutes into the amoxicillin pharmacokinetics experiment in Figure 6c,d. Plots show dynamics of green fluorescence (due to flowed fluorescein) of the highlighted feeding lane and individual trench sections from 0 to 600 minutes.

**Figure S10:**
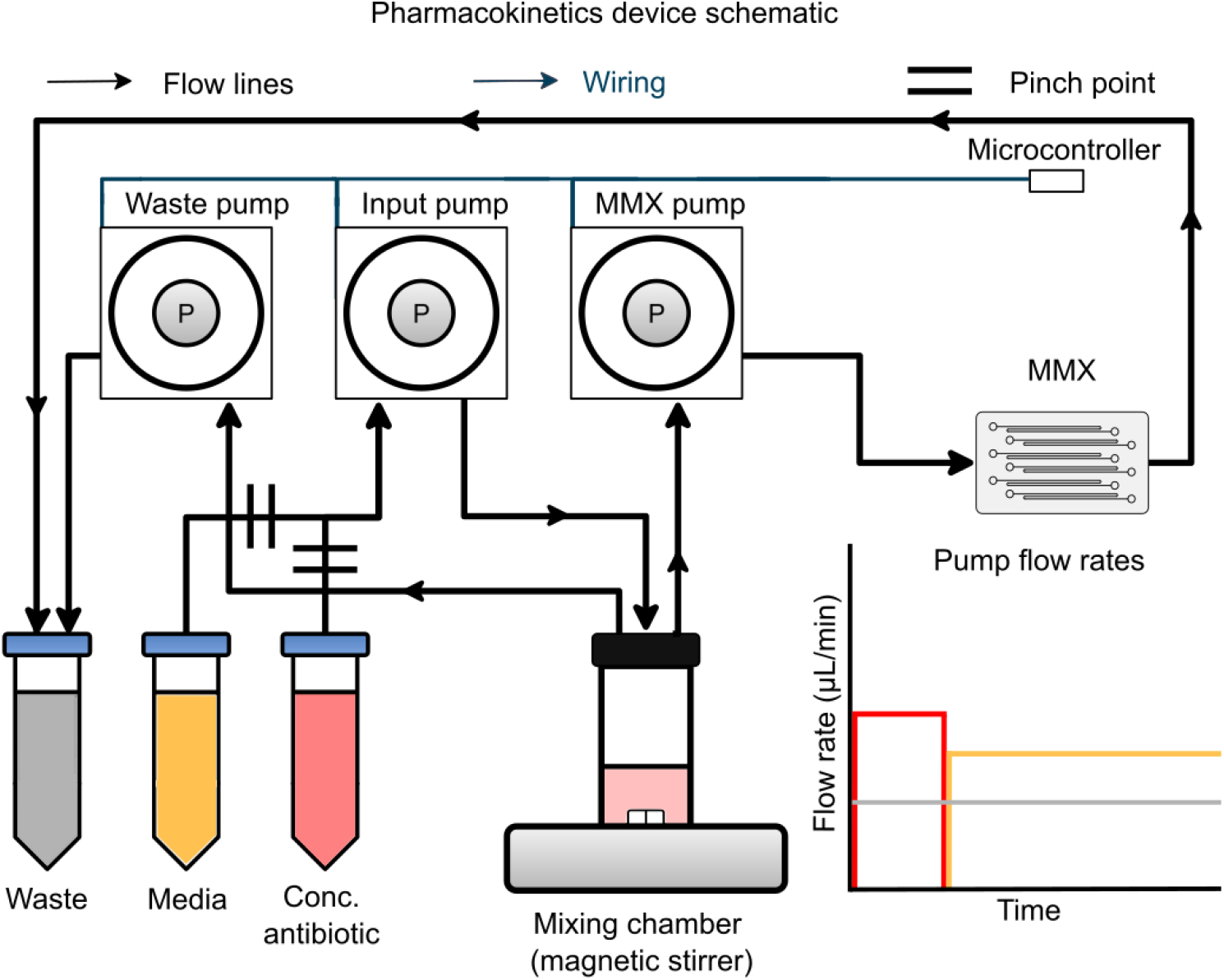
Schematic of the antibiotic pharmacokinetics system. Schematic depicting the multi-pump system for mimicking clinical antibiotic pharmacokinetics within the MMX device. Black lines indicate the direction of liquid flow from a tube containing concentrated antibiotic or from a tube containing fresh supplemented M9 media into a mixing chamber on a magnetic stirrer, which then flows into a MMX device loaded with cells and out to waste. Flow from either the concentrated antibiotic tube or fresh media tubes is controlled by an electronic pinch valve that swaps liquid flow between either concentrated antibiotic or fresh media being added to the mixing chamber.

### Supplementary Tables

**Table S1:** Minimum inhibitory concentration (MIC) of the antibiotics used in this study. *E. coli* MG1655 CGSC 6300 (“SB1”) were treated with prepared antibiotic stocks by the broth microdilution method performed as described in the methods section.

| Antibiotic | MIC ($\mu\text{g}/\text{mL}$) |
| --- | --- |
| Cefalexin | 8 |
| Ceftriaxone | 0.015625 |
| Amoxicillin | 4 |
| Ampicillin | 16 |

### Supplementary Videos

**Video 1**. Time-lapse imaging over a 30-minute period comparing loading dynamics in a MMX-design trench design (right), a MMX-design trench design with a smaller top chamber (middle), and a conventional mother machine trench (left). The MMX enables rapid loading, which is facilitated by evaporatively driven flow, substantially improving efficiency and throughput relative to the conventional, diffusion-based loading approach.

**Video 2**. Trench images, segmented masks, and corresponding length traces of examples of the different cell classes in response to antibiotic treatment. Red shaded region indicates 4 hours of treatment with 50 µg/mL ampicillin treatment. Top: Susceptible cells rapidly increase in size and subsequently lyse due to the antibiotic. Middle: Transiently tolerant cells initially start to increase in size, then experience a transient growth slowdown that facilitates their survival during antibiotic treatment. These cells often filament to form long cells and subsequently resolve into multiple cells following treatment end. Bottom: Starvation-triggered persisters maintain a slow increase in size from the start of treatment, facilitating their survival during antibiotic treatment, until later rapidly increasing in size.

## Supplementary Notes on Methods and Models

### Supplementary Note 1: Hi-DFA analysis pipeline

Microscopy images were saved as nd2 files. The image data and metadata in the nd2 files were extracted, and the images registered using PyStackReg[72] to eliminate stage drift. The trenches within each image were then identified from spikes in the mean *x* and *y* intensity profiles of the phase contrast channel, and extracted to a Zarr array[27] using a modified version of PyMMM[66].

The Zarr array of individual trench images was then segmented with Omnipose[25], using a model trained initially on appropriately optimised synthetic trench images generated by SyMBac[26] and subsequently fine-tuned by retraining on manually curated data from the previous iteration of the model output. Prior to segmentation, trench images were upscaled 2× in each spatial dimension using bicubic interpolation, yielding images of typical size 280 × 48 pixels from the native 140 × 24 pixels. This upscaling reduces the effect of pixelation for the small cell sizes typical of stationary-phase and early-resuscitation cells. The segmentation model was trained on equivalently upscaled images. Omnipose was trained for 4000 epochs, using default settings from the paper[25]. Segmentation masks were also saved as Zarr arrays.

Following image segmentation, small segmentation artefacts were removed by applying a minimum size threshold for masks, just below the size of the smallest real cell masks. Additional filters were applied for a minimum solidity threshold (which quantifies how convex masks are, calculated by regionprops from scikit-image[73]) and minimum distance from the edges of the image (to exclude segmentation artefacts outside of the trench).

The filtered segmentation masks were used to extract quantitative image features of the mother cell, using the mask at the top of the corresponding trench images, using regionprops. Extracted features typically included area, length, width, image intensity, solidity. The extracted data from regionprops was stored in a pandas dataframe, saved as a pickle file.

The data from the pickle file was subsequently used for dynamic fate analysis. Details on the time and treatment condition for each trench were added as metadata to the corresponding cell IDs of the dataframe. To identify trenches that contained at least one cell at the start of the experiment, we used a mask threshold, such that the trench could not lack a mask in more than 10% of the first 20 frames. A similar threshold was used to identify trenches that contained a cell at the end of the experiment, to minimise including segmentation artefacts.

The time-series of the measured length for each cell (which is the primary value used for subsequent analysis) then went through several cleanup steps. These cleanup steps included setting minimum length thresholds (to remove impossibly small cells, i.e. segmentation artefacts) and identifying positive and negative spikes where the length significantly increased/decreased then quickly reverted.

After these initial cleanup steps, division events were identified using SciPy[74] find_peaks function on the negative first derivative of the cleaned length data against time. A minimum prominence value was applied for identifying sufficiently large peaks (whilst excluding as many artefact peaks as possible), a minimum time between division events and minimum and maximum division event sizes were also applied to further minimise incorrectly identified division events.

To minimise segmentation noise in the length traces affecting calculated growth rates, the cleaned length trace data was smoothed using LOWESS smoothing (using a window size fraction set to a fixed number of timepoints, typically those comprising the surrounding ∼20 minutes of timepoints). Cell length drops owing to division events were temporarily “removed” to bring the divided cell lengths up to their pre-division lengths, then LOWESS smoothing was applied, before “adding back” the division events to the smoothed length traces.

Similar data cleanup parameters were used for all experiments, with the exception of the supra-MIC experiment which had 70 second frame intervals, instead of the typical 3 minute frame intervals, requiring appropriate modifications.

Growth rate was then calculated as the first derivative of the smoothed cell length traces and similar LOWESS smoothing was also applied to the growth-rate timeseries. The second derivative of the length trace was then calculated by taking the derivative of the growth rate trace, and light LOWESS smoothing applied. Growth slowdown events were identified from the second derivative of cell length timeseries. A slowdown was defined as a sustained drop, when the derivative remained below 0 (i.e. growth rate is slowing down) for above a threshold amount (defined by the integrated area above the curve). The growth slowdown was identified as terminated when the second derivative subsequently remained above 0 for a threshold amount (defined by the area below the curve exceeding a threshold amount relative to the identified accumulated negative area above the curve).

Cell survival was defined by the occurrence of a threshold number of division events following antibiotic treatment, typically assessed during the final 6 hours time window of the experiment. This threshold number of division events was chosen to be fairly generous, to also allow identification of slow growing cells or occasional missed division events during poorly segmented stretches. A filter for maximum % of acceptable NaN timepoints was also applied to minimise division event artefacts being included.

Following identification of cells surviving antibiotic treatment, to minimise the artefacts included in subsequent analysis, the trench images, corresponding masks, and corresponding growth traces were typically opened for each “survivor” in Napari[30], and Napari hotkeys were used to quickly classify ‘survivors’ as real or artefacts; this includes discarding cells in the rare case where the mother cell at the top of the trench dies, and one of the cells below the mother cell replaces it. During this manual data cleanup step, we typically also visually identified and classified the slow-growing starvation-triggered persisters, using a different hotkey. We note that based on the manually classified starvation-triggered persisters, an automatic classification criteria for this class of a mean growth rate below 0.0021 min^−1^ (∼10% of the maximum growth rate) for the first 120 minutes after introducing fresh media, and also for the first 60 minutes after introduction of fresh media (to exclude transient tolerant survivors), matches the manual classification quite well.

Survivors that were not classified as starvation-triggered persisters were then classified as either transiently tolerant survivors or exposure-limited survivors, based on the presence/absence of a significant growth slowdown event prior to the end of antibiotic treatment. Transiently tolerant survival events were identified by the presence of a slowdown event during antibiotic treatment that increased the integrated lost time of the cell from growth slowdown (using a similar definition to[71]) by at least 10 minutes.

After identifying cell classes, plots of cell length traces against time were created to visually confirm that everything seemed correct. Occasionally, a small number of “exposure-limited” traces that visibly showed growth slowdown, but were not classified as having significant slowdown events (usually due to segmentation errors), were manually classified into the transiently tolerant class.

After confirming cell classes, confidence intervals were calculated by bootstrapping with *n* = 10,000. These bootstrapped mean values and 95% confidence intervals were then used for subsequent analysis between different treatment conditions. Calculated growth rates were used for comparison between cell classes, or treatment conditions. To ensure that mean growth rate traces/heatmaps were minimally affected by artefacts, a small number of growth rate traces that clearly didn’t fit the raw data well (usually due to the raw data being messy, owing to occasional messy segmentation), were manually excluded from these analyses.

The Hi-DFA pipeline is available at https://github.com/kieranrabbott/Hi-DFA.

### Supplementary Note 2: Antibiotic pharmacokinetics system design and assembly

#### Hardware Configuration of the PK simulator

To simulate clinically relevant pharmacokinetic (PK) profiles in the microfluidic experiments, we developed a programmable flow-based system comprising three peristaltic pumps (Longer T100 & WX10 OEM), a mixing chamber, and a set of solenoid pinch-valve-controlled inlets for precise modulation of antibiotic concentration over time (schematic of the setup shown in Figure S10). The configuration was designed to deliver continuously varying antibiotic concentrations to the microfluidic device (MMX) while maintaining constant flow and volume within the system.

Two pumps were configured as outlet pumps, one directed flow from the mixing chamber through the MMX device, and the second removed excess liquid to waste, thereby maintaining stable pressure and flow conditions. A third inlet pump supplied media to the mixing chamber, which received input from either a fresh (antibiotic-free) or a concentrated (antibiotic-containing) medium source. These two input lines were joined via a Y-junction upstream of the mixing chamber. Flow selection between the fresh and concentrated inputs was controlled by a solenoid pinch valve (Cole-Parmer VapLock), which switched the inlet state according to a programmed time sequence to reproduce desired concentration-time profiles.

The mixing chamber consisted of a 20 mL borosilicate glass vial fitted with a custom cap containing three 0.75 mm-radius ports for inlet, outlet, and waste connections. A small magnetic stir bar placed inside the vial ensured rapid and complete mixing of the incoming streams when coupled with an external magnetic stirrer. The chamber thus acted as a continuously stirred reservoir where the antibiotic concentration evolved dynamically based on the inlet flow composition and dilution rate, reproducing exponential decay or complex clinical PK profiles as required.

Fluidic connections were made using 3/32 inch ID tubing. The outlet tubing was positioned below a 5 mL liquid line inside the chamber to ensure continuous withdrawal of liquid and to maintain a constant working volume. Flowrates for the inlet and outlet pumps were calibrated to match, maintaining a steady-state flow through the device. The concentration trajectory within the mixing chamber was verified experimentally using fluorescence dyes under identical flow conditions.

#### *in vitro* PK Simulator Logic

The PK setup was designed to be as minimal and reproducible as possible (Figure 6a), based on first-order pharmacokinetics to model both the concentration build-up (step-up to *C*_max_) and dilution (step-down) or the half-life. Pump speeds in each regime are controlled by an ESP32 microcontroller that encodes the calculated flowrates and their previously chosen time constants.

For example, for amoxicillin requiring approximately ∼80 minutes to achieve *C*_max_, the relationship between concentration and volume is given by:

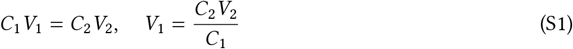

Substituting *C*_2_ (desired conc.) 2.5 mmol, *V*_2_ (initial volume) 5 mL, *C*_1_ (initial conc.):

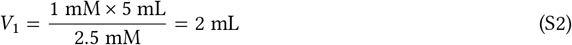

#### Flowrate Determination

The flowrate required to replace this calculated volume and achieve the desired concentration within the targeted time (*t*) is given by:

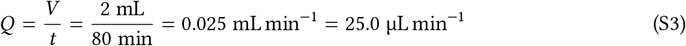

Where *Q* is the flowrate, maintaining constant volume requires:

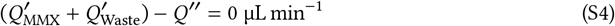

where *Q*^′^ is the output media and *Q*^′′^ is the input media (to the mixing chamber);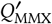 was fixed at 10 µL min^−1^.

#### Half-life (*t*_1/2_) flow profile

For calibration and validation of the pharmacokinetic (PK) setup, fluorescein was used as a non-reactive fluorescent tracer to quantify the generated concentration-time profiles of the antibiotics. The fluorescein intensity in the MMX device was continuously measured by fluorescence imaging to confirm that the system reproduced the intended temporal decay and steady-state characteristics under each programmed condition. In the half-life simulation phase, the objective was to achieve a dilution time constant equivalent to the reported pharmacokinetic half-life (*t*_1/2_) of the tested antibiotic. Here, *t*_1/2_ was defined as the time required to replace half of the total chamber volume (*V* ) with fresh medium. Using the flow-volume relationship the system’s embedded ESP32 microcontroller dynamically adjusted the pulse-width modulation (PWM) speed of each peristaltic pump to reproduce a first-order exponential decay of concentration over time. This enabled precise control of the antibiotic exposure profile, ensuring that the concentration delivered to the microfluidic device closely matched clinically relevant elimination kinetics. The ESP32 adjusted each pump’s PWM speed according to the required first-order decay profile.

#### Flowrate Calibration

Before each experiment, a series of flowrate calibration runs are completed, in which a Sensirion SLF3S-0600F flowrate monitor is connected to each fluidic line’s flow output and is monitored mapping PWM (Pulse Width Modulation) to µL min^−1^. The flowrate of which is recorded and PWM is tuned toward the desired value with a grid-search algorithm. Each pump is done accordingly, until the waste and fresh pumps are within ± 1 µL min^−1^ and the MMX pump is within ± 0.3–0.5 µL min^−1^. Sterilization is then conducted on the system as outlined in the methodology section and spent media is flowed through the system and the mixing chamber is filled with fresh M9 media.

### Supplementary Note 3: Mathematical Model of Transient Tolerance

## 1 General Model Framework

### 1.1 States and transitions

We consider a population of *N*_0_ cells at *t* = 0, the onset of antibiotic treatment and growth media replenishment, which can trigger exit from dormancy, at constant concentration *C* for duration *τ*. Each cell can occupy one of three states: dormant (*D*, not yet exited stationary phase), susceptible (*S*, actively growing and drug-susceptible), or transiently tolerant (*T*, actively growing with reduced drug susceptibility). Cells that die from either *S* or *T* enter an absorbing dead state. The correspondence between these model states and the experimentally observed post-treatment survivor classes is summarised in Table S2.

**Table S2:** Mapping between experimental survivor classes and model states. The three post-exposure survivor classes—persisters (*P* ⊂ *D*), exposure-limited survivors (*X* ⊂ *S*), and transiently tolerant survivors (*T* )—emerge naturally from the three model states *D, S*, and *T*, respectively (Figure 7c).

| Experimental term | Definition | Model state |
| --- | --- | --- |
| Lag-based survivor (incl. starvation-triggered persister, $P \subset D$) | Does not exit dormancy within $[0, \tau]$ ($L > \tau$); the extreme tail corresponds to classical starvation-triggered persisters | $D$ |
| Exposure-limited survivor ($X \subset S$) | Wakes under drug, remains in $S$ without switching, survives by timing (short $\tau$ or low $C$) | $S$ |
| Transiently tolerant survivor ($T$) | Wakes, transiently reduces susceptibility, survives with reduced but non-zero death rate | $T$ |

The population dynamics during the treatment phase (we consider transitions only during antibiotic exposure, not during post-treatment regrowth; see §3 for regrowth dynamics) are governed by

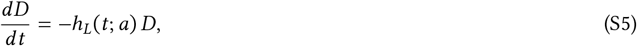

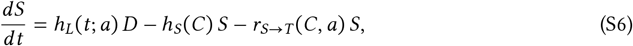

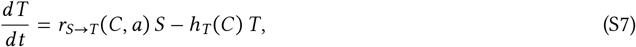

with initial conditions *D*(0) = *N*_0_, *S*(0) = *T* (0) = 0, and where:

- *h*_*L*_(*t*; *a*): instantaneous hazard of exiting dormancy at time *t*, given culture age *a*;
- *h*_*S*_(*C*): death rate of susceptible cells;
- *h*_*T*_ (*C*): death rate of transiently tolerant cells, with 0 < *h*_*T*_ (*C*) < *h*_*S*_(*C*) for all *C* > 0;
- *r*_*S*→*T*_ (*C, a*): rate of switching from *S* to *T* .

We impose no back-transition *T* → *S*, as we only consider transitions during the treatment phase, and this is consistent with the observation that cells entering the transiently tolerant state do not revert to full susceptibility during the treatment window, but rather die at their reduced growth rate as compared to fully susceptible cells.

### 1.2 Lag-time distribution

Individual lag times *L* follow a two-component mixture of an Erlang distribution (weight *w*, shape *k*, rate *λ*(*a*)) and a slow exponential (weight 1 − *w*, rate *λ*_*s*_):

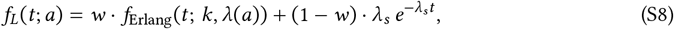

where the Erlang component has PDF, CDF, and survival function

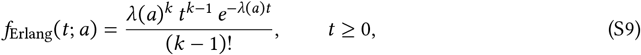

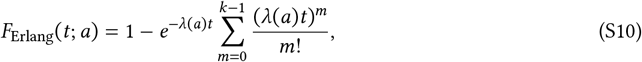

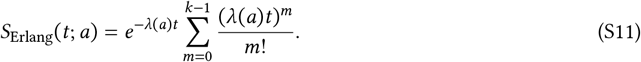

The overall survival function for the hybrid mixture is

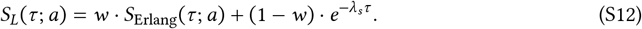

The mean lag time of the Erlang component is *μ*_lag_(*a*) = *k*/*λ*(*a*), modelled as a linear function of culture age:

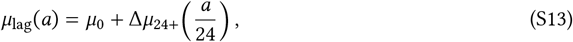

where *μ*_0_ is the baseline mean lag for a fresh culture and Δ*μ*_24+_ is the additional mean lag accumulated per 24 h of stationary phase. This gives *λ*(*a*) = *k*/*μ*_lag_(*a*).

The overall mean lag time under the mixture is 𝔼[*L*] = *w* · *μ*_lag_(*a*) + (1 − *w*)/*λ*_*s*_, and we define the *effective lag rate*

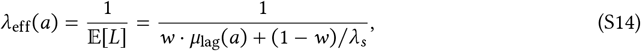

which replaces the bare Erlang rate *λ*(*a*) in all constraint penalties (§5). Unlike the Erlang rate *λ*(*a*), the slow-exit rate *λ*_*s*_ is modelled as age-independent: the deeply dormant subpopulation captured by this component is assumed to have entered a state whose exit kinetics are insensitive to the additional starvation stress accumulated over the culture ages tested. When *w* ≈ 0, the Erlang component is negligible and *λ*_eff_ ≈ *λ*_*s*_; when *w* = 1, *λ*_eff_ = *λ*(*a*) = *k*/*μ*_lag_(*a*).

The Erlang shape parameter *k* (selected by grid search over *k* = 2, …, 14) controls the regularity of wake times in the main component: larger *k* produces a sharper distribution around the mean (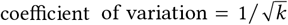). The slow-exponential component (rate *λ*_s_ ) captures a subpopulation of cells with extended lag times that are not well described by the Erlang tail; this heavy-tailed subpopulation corresponds to the slow phase of classical biphasic killing and includes cells that would conventionally be classified as starvation-triggered persisters.

### 1.3 Hazard functions

The concentration-dependent transition rates are expressed in terms of generic antibiotic pressure functions *φ*(*C*) that are monotonically increasing with *φ*(0) = 0:

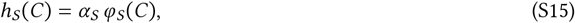

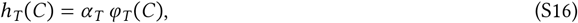

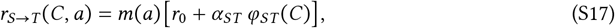

where *α*_*S*_, *α*_*T*_, *α*_*ST*_ are rate constants and *r*_0_ > 0 is a baseline switching rate that accounts for transient growth-slowdown events observed even in untreated resuscitating cells (Figure S5). The stress-history function *m*(*a*) = *a*/(*a* + *a*_50_) with *m* ∈ [0, 1) increases monotonically with culture age, capturing the observation that older cultures produce more transiently tolerant survivors. We impose *α*_*S*_ > *α*_*T*_ > 0 so that *h*_*S*_(*C*) > *h*_*T*_ (*C*) (for our choices of *h*_*S*_(*C*), *h*_*T*_ (*C*) described below) for all *C* > 0 when *φ*_*S*_ = *φ*_*T*_ .

## 2 Analytical Survivor Fractions

### 2.1 Convolution structure

A key property of the system (S5)–(S7) is that *h*_*S*_, *h*_*T*_, and *r*_*S*→*T*_ are constant during the treatment window (constant *C* and *a*). Only the dormancy-exit hazard *h*_*L*_(*t*; *a*) varies with time, so the *S*/*T* subsystem is a linear, time-invariant system driven by the time-varying inflow from *D*.

Solving (S5) gives *D*(*t*) = *N*_0_ *S*_*L*_(*t*; *a*), so the inflow rate into *S* at time *t* is *N*_0_*f*_*L*_(*t*; *a*). The surviving populations in *S* and *T* at time *t* are therefore convolutions:

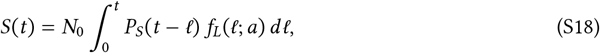

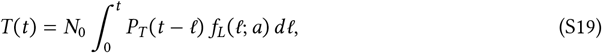

where *P*_*S*_(Δ) and *P*_*T*_ (Δ) are single-cell survival kernels for a cell exposed for duration Δ, derived below.

### 2.2 Derivation of the survival kernels

Consider a cell entering state *S* at time *ℓ*, subsequently exposed to antibiotic for duration Δ = *τ* − *ℓ*. Define the total exit rate from *S*:

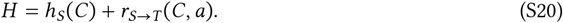

#### Kernel *P*_*S*_(Δ): surviving in *S*

The probability of remaining in *S* (neither dying nor switching) satisfies

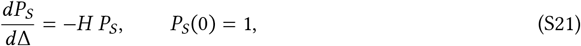

giving

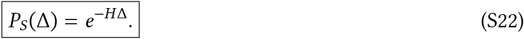

#### Kernel *P*_*T*_ (Δ): switching to *T* and surviving

A cell becomes a *T* -survivor by first switching *S* → *T* at some time *t* ∈ [0, Δ], then surviving in *T* for the remaining Δ − *t*. The probability density for the switch occurring at time *t*, given that the cell is still in *S* at time *t*, is

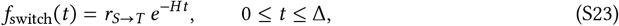

since *e*^−*Ht*^ is the probability of no event (neither death nor switch) in *S* up to *t*, and *r*_*S*→*T*_ is the instantaneous switching rate. Conditional on switching at *t*, survival in *T* for the remaining Δ − *t* is *e*^−*h*^*T* ^(Δ−*t*)^. Integrating over all possible switch times:

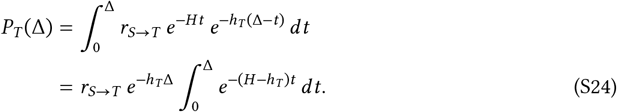

Evaluating the integral for the generic case *H* ≠ *h*_*T*_ and the degenerate case *H* = *h*_*T*_ :

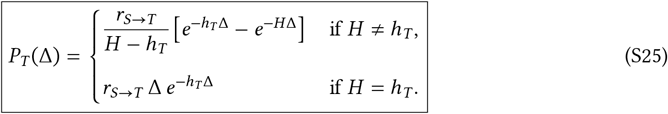

The remaining probability 1 − *P*_*S*_(Δ) − *P*_*T*_ (Δ) corresponds to death during exposure.

### 2.3 Population fractions at end of treatment

At time *τ*, the population is partitioned into four classes (Figure 7c):

#### Lag-based survivors (*P* ⊂ *D*)

Cells that have not yet resuscitated by the end of treatment (*L* > *τ*), the extreme tail of which corresponds to classical starvation-triggered persisters:

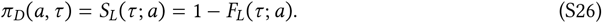

#### Exposure-limited survivors (*X* ⊂ *S*)

Cells that resuscitated during treatment (*L* ≤ *τ*), entered *S*, and survived in *S* until *τ* without switching to *T* . These are not a mechanistically distinct class, but rather cells whose growth trajectory and treatment timing allowed them to escape killing:

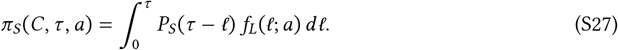

#### Transiently tolerant survivors (*T* )

Cells that resuscitated, switched to *T*, and survived in *T* until *τ*:

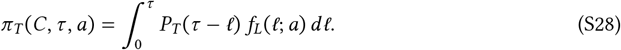

#### Dead

π_dead_ = 1 − π_*D*_ − π_*S*_ − π_*T*_ .

*Remark*. These expressions are unconditional population fractions (over the entire initial population). They can equivalently be derived by writing the conditional survivor fractions among resuscitating cells (those with *L* ≤ *τ*) and multiplying by the resuscitation probability *F*_*L*_(*τ*; *a*); the prefactors cancel.

### 2.4 Closed-form evaluation

Define the *discounted wake integral*

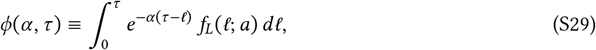

which has a direct physical interpretation: *ϕ*(*α, τ*) is the total surviving mass of cells that woke during [0, *τ*] and experienced a constant-rate clearance process with rate *α* for their remaining exposure time. The survivor fractions then take the compact forms

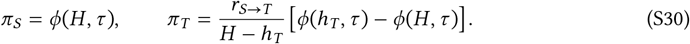

For the hybrid lag distribution (S8), *ϕ* decomposes as a weighted sum:

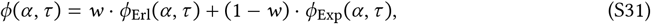

where the Erlang component is evaluated via the *J*_*m*_ recurrence:

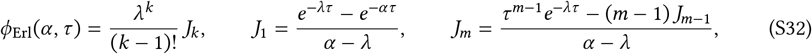

and the exponential component has the closed form

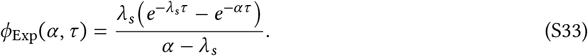

When *α* ≈ *λ* or *α* ≈ *λ*_*s*_, Taylor expansions are used to avoid catastrophic cancellation (threshold |Δ|*τ* < 10^−8^). In practice, the product *e*^−*ατ*^ · *J*_*k*_ is evaluated by carrying the exponential factor into the recurrence, operating on the numerically stable difference *e*^−*λτ*^ − *e*^−*ατ*^.

## 3 Post-Treatment Regrowth Dynamics

### 3.1 Setup

At *t* = *τ* the antibiotic is removed. Let *u* = *t* −*τ* ≥ 0 denote the time since drug removal. Both transiently tolerant and exposure-limited survivors have already exited dormancy during treatment; thus, upon drug removal, transiently tolerant cells are assumed to revert immediately to a growing phenotype, so that all awakened survivors resume exponential growth immediately (residual lag *L*_res_ = 0):

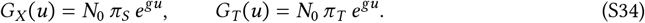

Lag-based survivors (including persisters, *P* ⊂ *D*), by contrast, retain a residual lag of *L*_res_ = *L* − *τ*: they must first complete the remaining stages of their dormancy exit before contributing to the growing population. This delay is the key asymmetry: transiently tolerant and exposure-limited survivors have a head start in seeding regrowth.

### 3.2 Residual lag via the linear-chain trick

The Erlang component of the hybrid lag distribution (weight *w*, shape *k*) can be represented as the sum of *k* independent exponential stages, each with rate *λ*. The exponential component (weight 1 − *w*, rate *λ*_*s*_) contributes a single additional stage. At *u* = 0, the surviving dormant mass *N*_0_π_*D*_ is distributed across these stages according to the hybrid survival function (S12).

For the Erlang component, the linear-chain dynamics are:

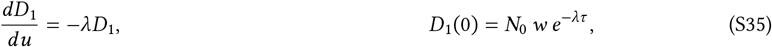

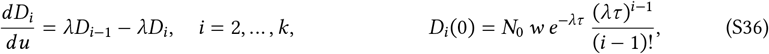

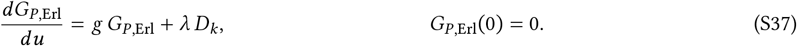

For the exponential component:

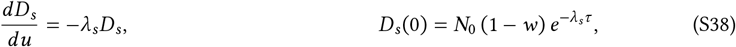

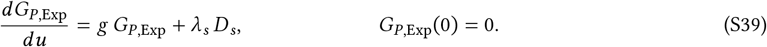

The total growing pool from persisters is *G*_*P*_ = *G*_*P*,Erl_ + *G*_*P*,Exp_. One can verify that the initial masses sum correctly: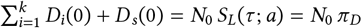.

### 3.3 Dormant descendant contribution

By variation of parameters:

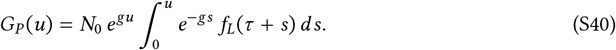

This integral accounts for the fact that lag-based survivors (including persisters) that complete their residual lag at post-treatment time *s* contribute to the growing population, but their descendants are discounted by the factor *e*^−*gs*^ relative to cells that were already growing from *u* = 0. For the hybrid lag distribution (S8), both the Erlang and exponential components contribute independently to this integral.

### 3.4 Descendant shares

The total regrowing population is *N* (*u*) = *G*_*X*_ (*u*) + *G*_*T*_ (*u*) + *G*_*P*_ (*u*). The fractional contribution of each survivor class is

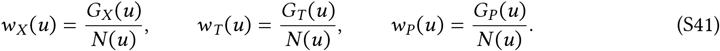

Cancelling the common factor *N*_0_*e*^*gu*^:

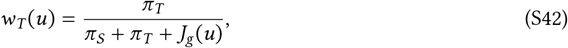

where

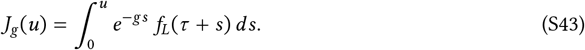

Since *J*_*g*_ (0) = 0 and *J*_*g*_ (*u*) increases monotonically to its asymptotic value *J*_*g*_ (∞), the transiently tolerant share *w*_*T*_ (*u*) is maximal immediately after drug removal and decreases as lag-based survivors (persisters) progressively complete their residual lag and enter the growing population. The structure of Equation (S42) makes explicit that transiently tolerant-descended cells dominate early regrowth whenever π_*T*_ is comparable to or exceeds π_*S*_, because persisters (*P* ⊂ *D*) contribute only with a delay penalised by their residual lag *L*_res_ = *L* − *τ*.

For the hybrid lag distribution, *J*_*g*_ (*u*) decomposes into Erlang and exponential contributions:

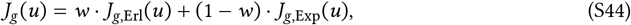

where the Erlang component is evaluated using the regularised incomplete gamma function with *α* = *λ* + *g*:

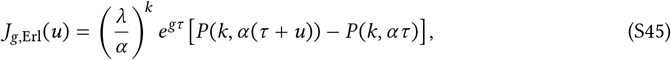

where *P* (*k, x*) = *γ*_reg_(*k, x*) is the regularised lower incomplete gamma function, and the exponential component has the closed form

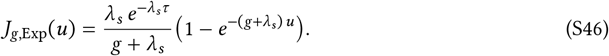

Both components permit closed-form solutions: the Erlang component because *k* is an integer, and the exponential component by direct integration.

## 4 Model hazard functions

The primary model uses Hill functions for the antibiotic pressure:

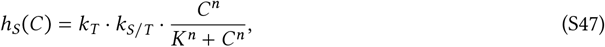

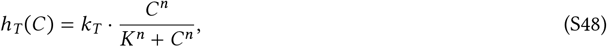

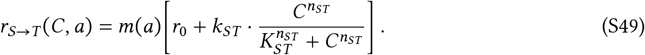

The death hazards *h*_*S*_ and *h*_*T*_ share a common Hill coefficient *n* and half-maximal concentration *K*, differing only in their maximal rates: *α*_*S*_ = *k*_*T*_ · *k*_*S*/*T*_ for susceptible cells and *α*_*T*_ = *k*_*T*_ for transiently tolerant cells. The constraint *k*_*S*/*T*_ > 1 ensures *h*_*S*_(*C*) > *h*_*T*_ (*C*) for all *C* > 0. Sharing *n* and *K* encodes the assumption that transient tolerance acts by reducing the maximal kill rate rather than shifting the dose–response curve, and reduces the number of free parameters.

### Wake-rate constraint

We impose *λ*_eff_(*a*) ≤ *k*_*T*_ for all culture ages in the dataset, where *λ*_eff_(*a*) is the effective lag rate defined in (S14). Since *h*_*T*_ (*C*) → *k*_*T*_ as *C* → ∞, this constraint ensures that at saturating antibiotic concentrations, the transiently tolerant kill rate equals or exceeds the effective rate at which dormant cells wake into the susceptible pool. Without this constraint, the model could predict that *T* cells are generated (via *S* → *T* switching) faster than they are cleared at arbitrarily high concentrations, implying implausible persistence of the transiently tolerant class under conditions where we experimentally observe near-complete killing (Figure 4e, 250 µg/mL). With the constraint in place, high *C* and long *τ* efficiently clear both *S* and *T* cells, and the survivor pool contracts to the residual tail π_*D*_, which itself becomes vanishingly small at treatment durations well beyond the mean lag time. The constraint is implemented as a quadratic penalty (see §5). Using *λ*_eff_ rather than the bare Erlang rate *λ* is critical when the mixture weight *w* ≈ 0: in that regime, the Erlang rate can be very large while the actual wake rate is governed by the slow exponential component *λ*_*s*_.

Two variants of the Hill model are compared:

1. **Hill-Hybrid** (primary, 13 free parameters): uses the full hybrid Erlang+Exponential lag distribution (S8) with both *w* and *λ*_*s*_ fitted. This is the production model (Table S3).
2. **Hill-Erlang** (11 free parameters): uses a pure Erlang lag distribution (*w* = 1 fixed, *λ*_*s*_ not fitted). This intermediate model tests whether the slow-exponential lag component improves fit quality.
3. Additionally, a reduced **Linear minimal** model variant was also fitted to the data for model comparison purposes (§5.4): a **Linear-Hazard minimal model** (7 parameters; hazards proportional to *C* without saturation, pure Erlang lag). The Linear minimal model is thus low-concentration approximation of the Hill model.

**Table S3:** Model parameters and their presence across model variants.

| Symbol | Biological meaning | Fitted | Hill-Hybrid | Hill-Erlang | Linear |
| --- | --- | --- | --- | --- | --- |
| $k$ | Erlang shape (number of lag stages) | Yes | ✓ | ✓ | ✓ |
| $w$ | Erlang mixture weight | Yes | ✓ | – | – |
| $\lambda_s$ | Slow-exit rate, exponential component ($\text{h}^{-1}$) | Yes | ✓ | – | – |
| $\mu_0$ | Baseline mean lag time, Erlang component (h) | Yes | ✓ | ✓ | ✓ |
| $\Delta\mu_{24+}$ | Additional mean lag per 24 h of starvation (h) | Yes | ✓ | ✓ | ✓ |
| $k_T$ | Maximal kill rate, $T$ state ($\text{h}^{-1}$) | Yes | ✓ | ✓ | ✓ |
| $k_{S/T}$ | Kill-rate ratio $h_S/h_T$ ($> 1$) | Yes | ✓ | ✓ | ✓ |
| $k_{ST}$ | Maximal $S \rightarrow T$ switching rate ($\text{h}^{-1}$) | Yes | ✓ | ✓ | ✓ |
| $K$ | Shared half-max. conc. for $h_S, h_T$ ($\mu\text{g mL}^{-1}$) | Yes | ✓ | ✓ | – |
| $K_{ST}$ | Half-max. conc. for $r_{S \rightarrow T}$ ($\mu\text{g mL}^{-1}$) | Yes | ✓ | ✓ | – |
| $n$ | Shared Hill coefficient for $h_S, h_T$ | Yes | ✓ | ✓ | – |
| $n_{ST}$ | Hill coefficient for $r_{S \rightarrow T}$ | Yes | ✓ | ✓ | – |
| $a_{50}$ | Culture age at half-maximal stress history (h) | Yes | ✓ | ✓ | ✓ |
| $r_0$ | Baseline $S \rightarrow T$ switching rate at $C = 0$ ($\text{h}^{-1}$) | No | ✓ | ✓ | ✓ |
| $g$ | Post-treatment balanced growth rate ($\text{h}^{-1}$); not fitted, as it does not enter the survival fraction predictions | Fixed | ✓ | ✓ | ✓ |

## 5 Fitting Procedure and Sensitivity Analysis

### 5.1 Parameter estimation

All parameters were fitted by minimising a composite objective in log-space using L-BFGS-B (SciPy 1.14) with multistart optimisation (240 random initialisations, jitter *σ* = 3.5 in log-space). The composite objective is

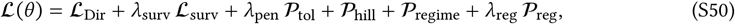

where the six terms are described below. The Erlang shape *k* was selected by grid search over *k* = 2, …, 14, refitting all continuous parameters at each value of *k*. Parameter bounds are listed in Table S4.

#### Four-class Dirichlet NLL (*ℒ* _Dir_)

For each experimental condition *j* with observed fractions 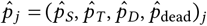 and predicted fractions *p*_*j*_ :

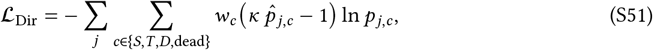

with Dirichlet concentration parameter *κ* = 5000 and optional class weights *w*_*c*_.

#### Survivor-composition NLL *(ℒ* _surv_)

The four-class Dirichlet NLL is dominated by the dead class at high concentrations (often >99% dead), leaving almost no gradient signal about *which state* the survivors belong to. The survivor-composition NLL addresses this by operating on the renormalised survivor fractions:

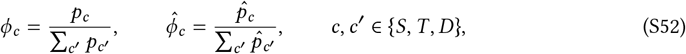

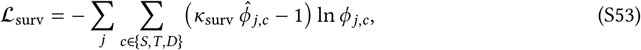

with *κ*_surv_ = 500. Conditions with zero observed survivors are excluded. This term ensures that the model correctly predicts *D*-dominance among survivors at high *C* even though the total surviving fraction is vanishingly small.

#### Tolerance penalty (*𝒫* _tol_)

Enforces *h*_*T*_ (*C*)/*h*_*S*_(*C*) < *ρ* (default *ρ* = 0.8) across a concentration grid *C* ∈ [0.05, 300] and representative ages, using a softplus barrier. With shared *K* and *n*, this ratio is the constant 1/*k*_*S*/*T*_, so the penalty reduces to enforcing *k*_*S*/*T*_ > 1/*ρ*.

#### Hill-lambda constraint (*𝒫* _hill_)

A quadratic penalty enforcing *λ*_eff_(*a*) ≤ *k*_*T*_ for all culture ages in the dataset:

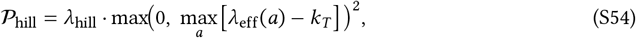

where *λ*_hill_ = 10^6^. This uses the effective lag rate *λ*_eff_(*a*) from (S14) rather than the bare Erlang rate, which is critical when *w* ≈ 0.

#### Regime-dominance penalty *(𝒫* _regime_)

Encourages *k*_*T*_ /*λ*_eff_(*a*) ≥ *β*_min_ (default *β*_min_ = 2), targeting the boundary where permanent *T* -over-*D* dominance becomes impossible at saturating *C*:

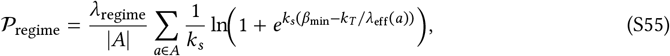

with *λ*_regime_ = 10^4^.

#### Parameter regularisation (*𝒫* _reg_)

Softplus penalties on biologically implausible parameter values: *n* > 10, *n*_*ST*_ > 10, *r*_0_ > 0.5, and *k*_*S*/*T*_ < 1.5.

The default hyperparameters are *λ*_surv_ = 1, *λ*_pen_ = 10^4^, and *λ*_reg_ = 10^3^.

**Table S4:**
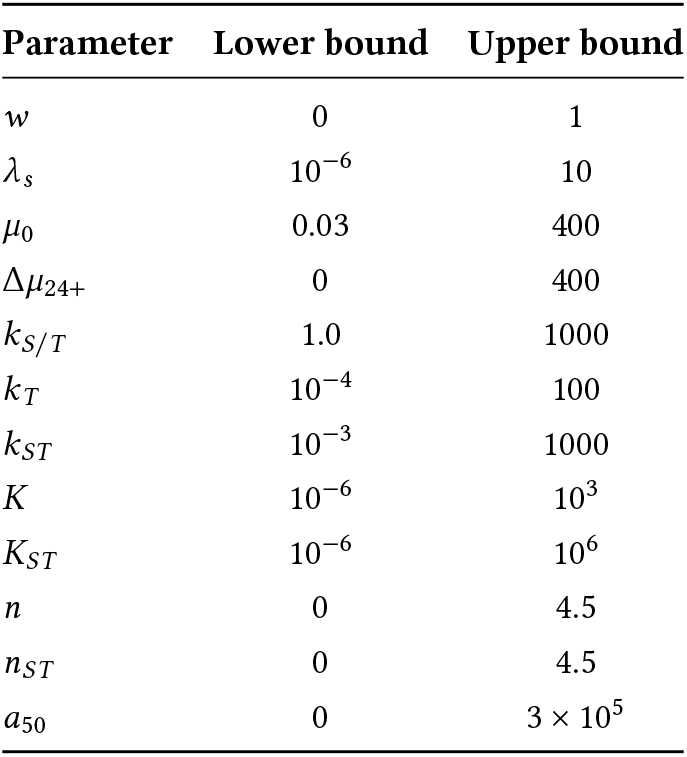
Parameter bounds (natural scale). The Hill-Hybrid model fits all 13 parameters; the Hill-Erlang model fixes *w* = 1 and omits *λ*_*s*_; the Linear model additionally omits *K, K*_*ST*_, *n, n*_*ST*_ and fixes *n* = *n*_*ST*_ = 1.

| Parameter | Lower bound | Upper bound |
| --- | --- | --- |
| $w$ | 0 | 1 |
| $\lambda_s$ | $10^{-6}$ | 10 |
| $\mu_0$ | 0.03 | 400 |
| $\Delta\mu_{24+}$ | 0 | 400 |
| $k_{S/T}$ | 1.0 | 1000 |
| $k_T$ | $10^{-4}$ | 100 |
| $k_{ST}$ | $10^{-3}$ | 1000 |
| $K$ | $10^{-6}$ | $10^3$ |
| $K_{ST}$ | $10^{-6}$ | $10^6$ |
| $n$ | 0 | 4.5 |
| $n_{ST}$ | 0 | 4.5 |
| $a_{50}$ | 0 | $3 \times 10^5$ |

**Table S5:** Maximum-likelihood parameter estimates for the Hill-Hybrid model (*k* = 6, 13 free parameters). The Erlang shape *k* was selected by grid search; all other parameters were fitted by L-BFGS-B multistart optimisation.

| Symbol | Parameter | MLE |
| --- | --- | --- |
| $k$ | Erlang shape | 6 |
| $w$ | Erlang mixture weight | 0.999 |
| $\lambda_s$ | Slow-exit rate ($\text{h}^{-1}$) | $1.63 \times 10^{-4}$ |
| $\mu_0$ | Baseline mean lag (h) | 0.715 |
| $\Delta\mu_{24+}$ | Additional mean lag per 24 h (h) | 0.184 |
| $k_T$ | Max. kill rate, $T$ ($\text{h}^{-1}$) | 3.55 |
| $k_{S/T}$ | Kill-rate ratio $h_S/h_T$ | 2.26 |
| $k_{ST}$ | Max. switching rate ($\text{h}^{-1}$) | 1.74 |
| $K$ | Half-max. conc., $h_S/h_T$ ($\mu\text{g mL}^{-1}$) | 65.7 |
| $K_{ST}$ | Half-max. conc., $r_{S \rightarrow T}$ ($\mu\text{g mL}^{-1}$) | 32.6 |
| $n$ | Hill coefficient, $h_S/h_T$ | 1.19 |
| $n_{ST}$ | Hill coefficient, $r_{S \rightarrow T}$ | 2.36 |
| $a_{50}$ | Half-max. stress history (h) | 26.0 |
| $r_0$ | Baseline switching rate ($\text{h}^{-1}$) | 0.0608 |

### 5.2 Sensitivity analysis

Parameter uncertainty was assessed by Laplace approximation at the MLE: the Hessian of the negative log-likelihood was computed by central finite differences, inverted to obtain an approximate covariance matrix in log-space, and used to generate 5 000 multivariate normal draws. These draws were propagated through the model to obtain 95% prediction intervals on all survivor fractions (reported on the predicted-versus-observed plots, Figure 8a–d). The resulting marginal posterior distributions for all 13 Hill-Hybrid parameters are shown in Figure S11.

To assess whether the predicted survivor composition is robust to parameter uncertainty, we propagated the posterior draws through the model. Posterior histograms of the survivor fractions *φ*_*T*_, *φ*_*S*_, and *φ*_*D*_ at six representative treatment conditions are shown in Figures S12-S14. The corresponding one-dimensional composition envelopes-MLE curves with 95% posterior bands for all three fractions as a function of concentration *C* and treatment duration *τ*—are shown in Figure S15.

**Figure S11:**
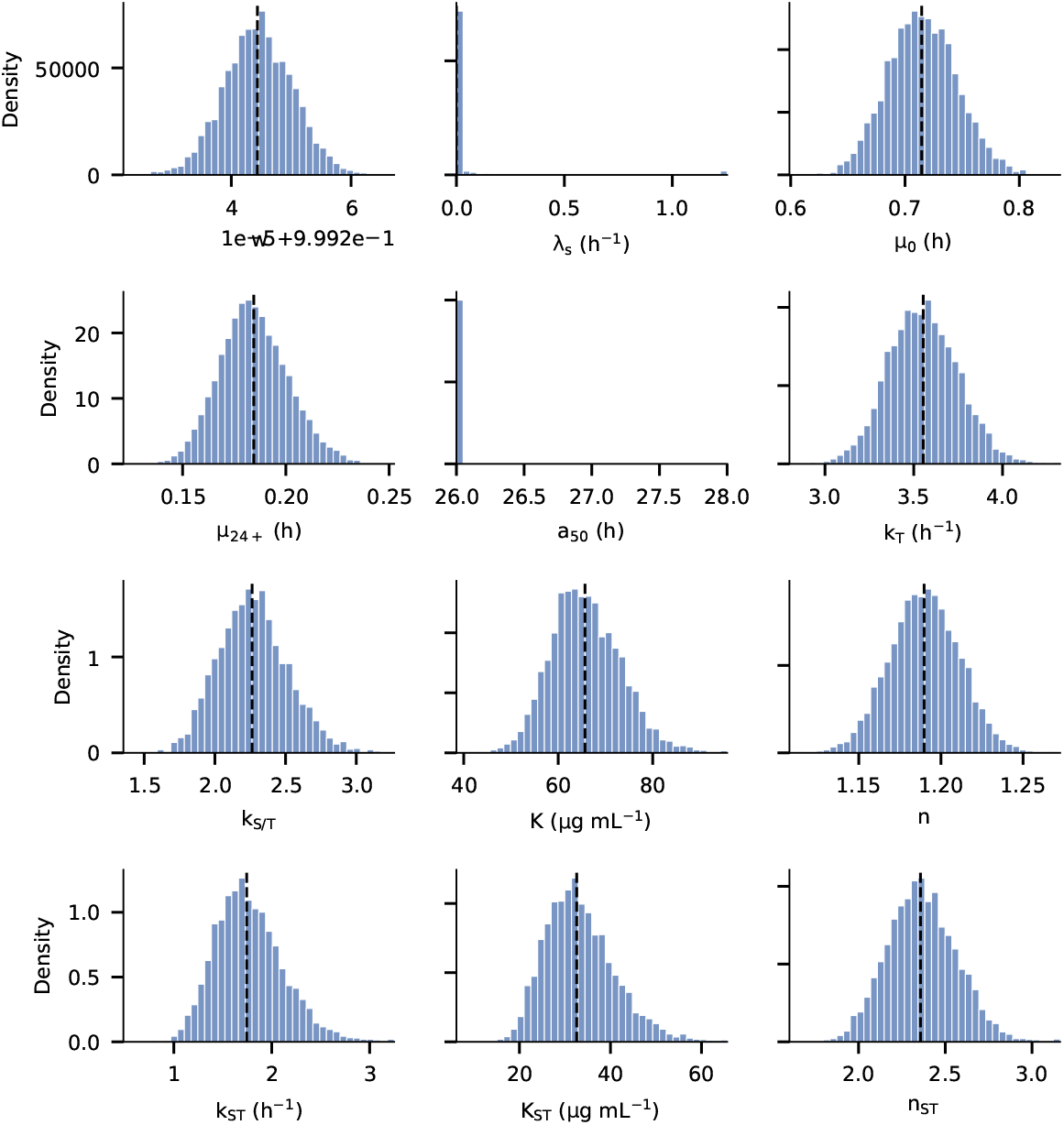
Marginal posterior distributions for all 13 Hill-Hybrid model parameters, obtained from the Laplace approximation (5 000 multivariate normal draws in log-space, transformed to natural scale). Dashed vertical lines indicate the maximum-likelihood estimates.

### 5.3 Fit quality for the lag-based survivor class

The model accurately reproduces the observed fractions of exposure-limited survivors (*X* ⊂ *S*), transiently tolerant survivors (*T* ), and dead cells across the full experimental range (Figure 8a–c). The lag-based survivor fraction (*P* ⊂ *D*, Figure 8d) shows a systematic underprediction, which we discuss here.

#### Origin of the deviation

The predicted lag-based survivor fraction π_*D*_(*τ, a*) = *S*_*L*_(*τ*; *a*) depends only on the lag-time distribution and the treatment duration; it is independent of antibiotic concentration (Equation S26). For a given (*τ, a*), the model therefore predicts a single value of π_*D*_ regardless of *C*. The observed fractions, however, represent at most 2–14 cells per condition (out of >10 000 cells per condition), placing them at the limit of reliable frequency estimation. Small-number fluctuations in this regime can produce apparent deviations of an order of magnitude or more. In addition, the Erlang distribution, while well suited to capturing the bulk of the resuscitation dynamics, has exponentially decaying tails. The real lag-time distribution likely has a heavier tail, consistent with the observation that a small number of cells with exceptionally long lag times persist even at *τ* = 8 h, where the Erlang survival function is negligible (*S*_*L*_(8; *a*=24) ≈ 10^−14^).

#### Classification boundary at short treatment durations

The model classifies any cell with *L* > *τ* as a lag-based survivor, without imposing a minimum lag duration for what constitutes a “true” persister. At short treatment durations (*τ* < ∼1 h), a substantial fraction of the population has lag times exceeding the brief treatment window—these are ordinary slow-waking cells, not the rare, deeply dormant persisters that survive prolonged high-dose exposure. This accounts for the model predicting a high lag-based survivor fraction at short *τ* across all concentrations (visible as the blue band in Figure 8f). Experimentally, subsequent growth dynamics distinguish these cells from deep persisters, but the model’s *L* > *τ* criterion does not make this distinction.

#### Impact on conclusions

The underprediction of the persister fraction does not affect the model’s central predictions for two reasons. First, the post-treatment regrowth analysis (§3) shows that persisters contribute to regrowth only after completing their residual lag (*L*_res_ = *L* − *τ*), which introduces a substantial delay relative to *T* and *X* ⊂ *S* survivors that resume growth immediately. Even if the true persister fraction were an order of magnitude higher than predicted, the growth-discount penalty (Equation S42) ensures that *T* -descended cells still dominate early regrowth under clinically relevant conditions. Second, the conditions where persisters are most underpredicted (high *C*, long *τ*) are precisely those where near-complete killing is observed (> 99.9% dead), so the absolute number of survivors of any class is very small.

### 5.4 Model comparison

The three model variants defined in §4 were fitted to the same dataset (*n* = 10 conditions) and compared using the four-class Dirichlet NLL (*κ* = 5000), AIC, and BIC (Table S6).

**Table S6:** Model comparison (lower is better for all criteria). Δ values are relative to the Hill-Hybrid model.

| Model | Free params ($k$) | NLL | AIC | BIC |
| --- | --- | --- | --- | --- |
| Hill-Hybrid (primary) | 13 | 15 208.6 | 30 443.1 | 30 447.1 |
| Hill-Erlang | 11 | 15 275.7 | 30 573.4 | 30 576.8 |
| Linear (minimal) | 7 | 15 957.5 | 31 929.1 | 31 931.2 |

Both AIC and BIC favour the Hill-Hybrid model, indicating that the additional parameters are supported by the data rather than reflecting overfitting. Critically, the central prediction, that transiently tolerant-descended cells dominate post-treatment regrowth under clinically relevant conditions, is reproduced by all three model variants, confirming that this conclusion is robust to model specification and does not depend on the specific parametric form of the hazard functions. The models diverge at high antibiotic concentrations (*C* ≥ 50 µg mL^−1^), where the reduced models cannot capture dose–response saturation, and in the persister tail, where the hybrid lag distribution better reproduces the observed frequency of late-waking cells.

**Figure S12:**
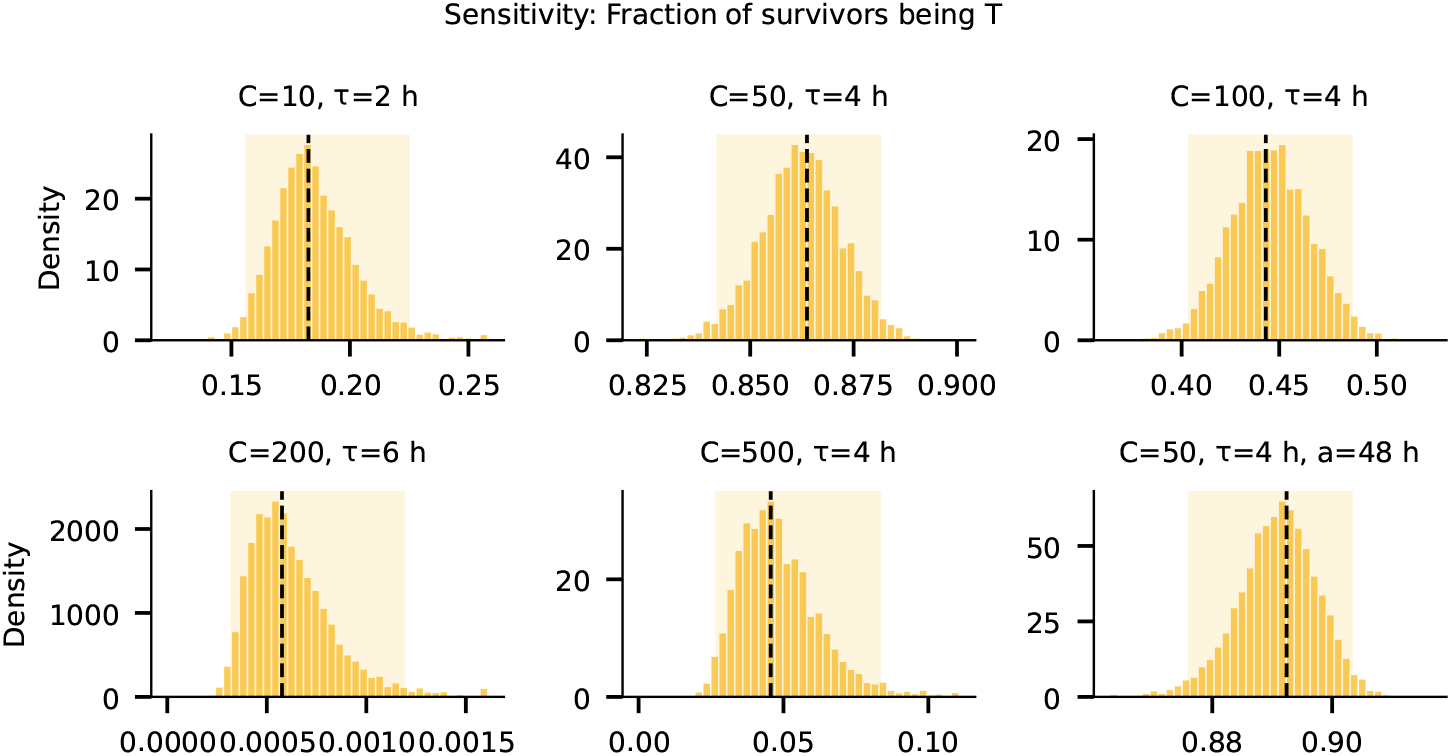
Sensitivity analysis: fraction of survivors being tolerant. Posterior histograms of the tolerant survivor fraction π_*T*_ /(π_*S*_ + π_*T*_ + π_*D*_) at six representative (*C, τ, a*) conditions, computed from 5 000 Laplace-approximation draws. Dashed lines indicate the MLE; shaded bands mark the 95% confidence interval.

**Figure S13:**
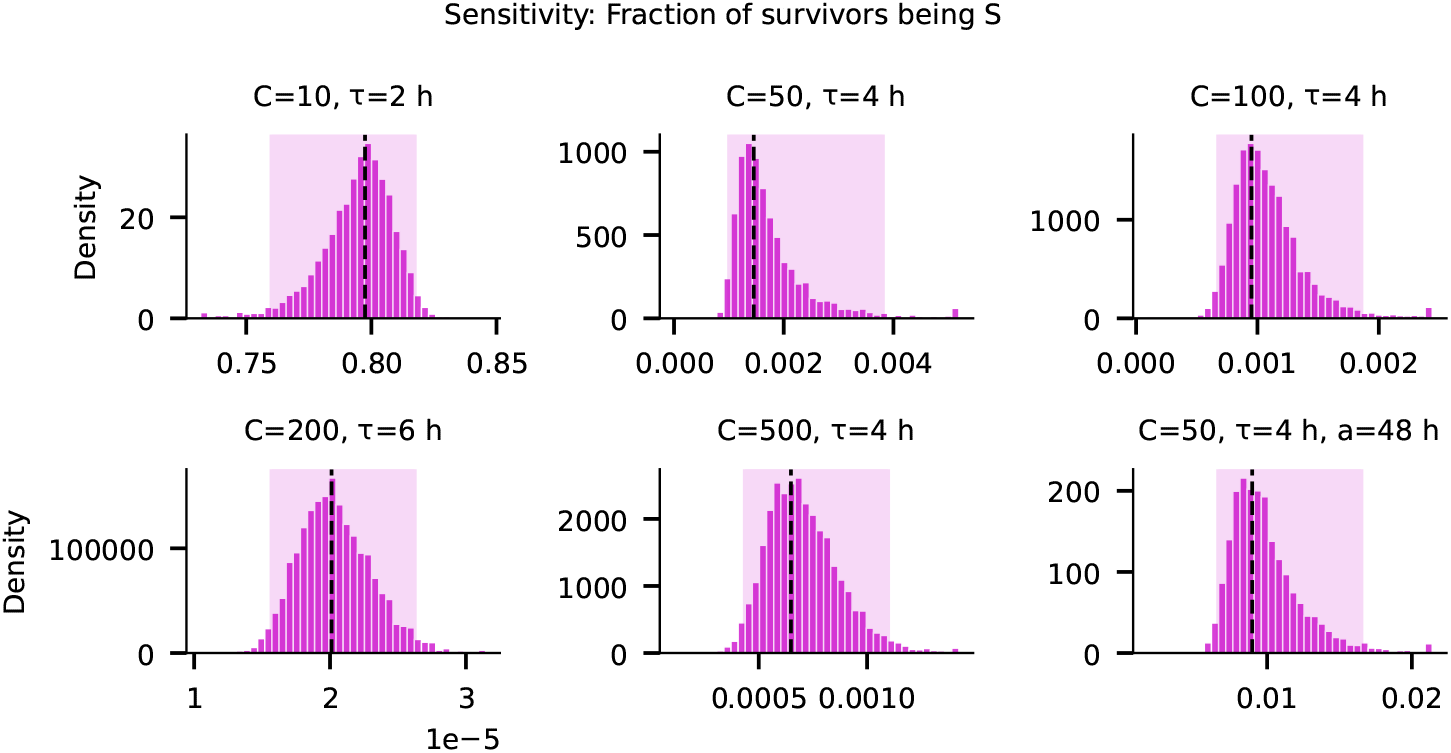
Sensitivity analysis: fraction of survivors being susceptible. Posterior histograms of the exposure-limited survivor fraction π_*S*_/(π_*S*_ + π_*T*_ + π_*D*_) at the same six conditions as Figure S12.

**Figure S14:**
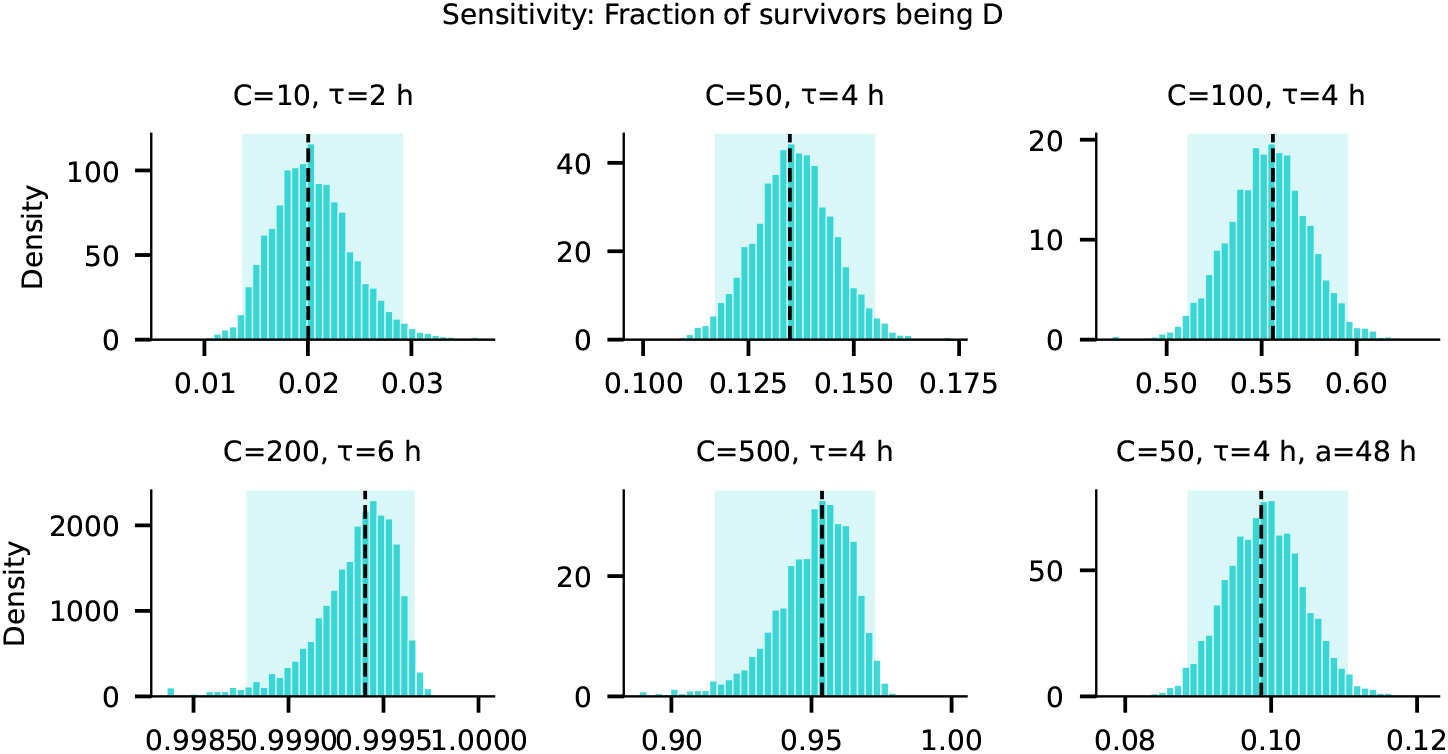
Sensitivity analysis: fraction of survivors being persisters. Posterior histograms of the lag-based (persister) survivor fraction π_*D*_/(π_*S*_ + π_*T*_ + π_*D*_) at the same six conditions as Figure S12.

**Figure S15:**
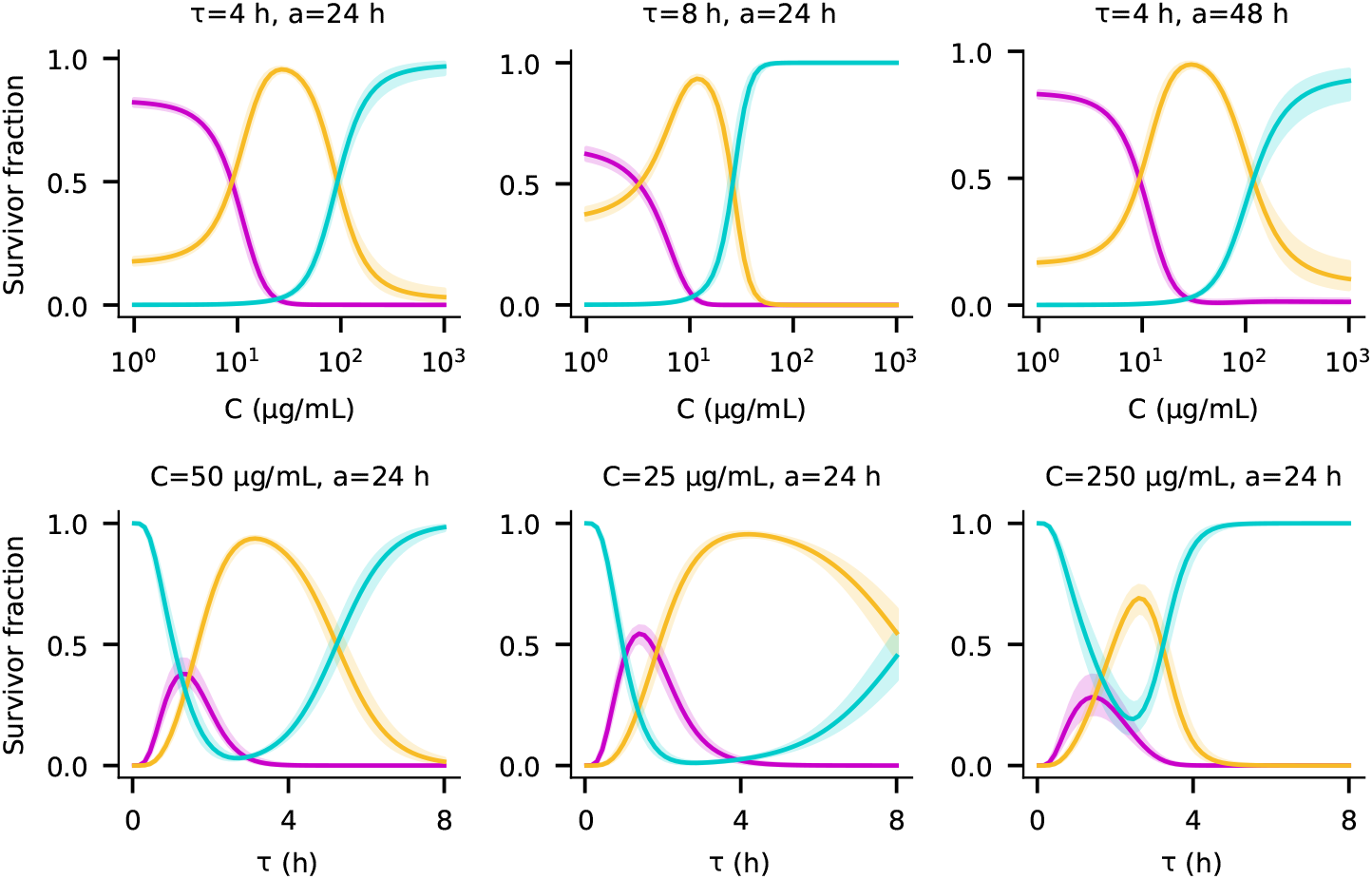
Posterior composition envelopes. MLE survivor-fraction curves (solid) with 95% posterior bands (shaded) for all three survivor classes. Top row: composition vs. antibiotic concentration *C* at three (*τ, a*) slices. Bottom row: composition vs. treatment duration *τ* at three (*C, a*) slices.

